# CytoAnalyst: A web-based platform for comprehensive single-cell RNA sequencing analysis

**DOI:** 10.1101/2025.04.14.647594

**Authors:** Phi Bya, Duy Tran, Khoi Nguyen, Sorin Draghici, Tin Nguyen

## Abstract

Single-cell technologies have revolutionized our ability to study cellular heterogeneity and dynamics at unprecedented resolutions. In this fast-growing field, it becomes increasingly challenging to navigate the vast amount of tools and steps for analysis. It is particularly difficult to integrate and analyze large datasets that require extensive collaborations and customized pipelines to obtain robust results. We present CytoAnalyst, a web-based platform that offers a number of important advantages over existing tools for single-cell analysis. First, the platform enables custom pipeline configuration using an efficient study management system and a broad range of analysis modules. Second, the platform supports parallel analysis instances, facilitating the comprehensive comparison of different methods or parameter settings available at each analysis step. Third, the advanced sharing system facilitates real-time synchronization among team members and seamless analysis continuation across different devices. Finally, the multi-grid visualization system supports simultaneous display of different data aspects, allowing for the comparison of multiple labels and plots side-by-side for comprehensive data insights, with the ability to save and reload visualization settings at any analysis step. The platform incorporates multiple blending modes, allowing users to combine plots in various ways for comprehensive data exploration. CytoAnalyst supports a high level of analytical rigor while providing user-friendly and flexible operations through its carefully designed interface and extensive documentation. The platform supports all major web browsers and is freely available at https://cytoanalyst.tinnguyen-lab.com.

## 1 Background

Single-cell RNA sequencing (scRNA-Seq) has emerged as a transformative technology in biomedical research, enabling unprecedented insights into cellular heterogeneity, rare cell populations, and dynamic biological processes [1]. Single-cell analysis often follows a complex workflow that involves multiple steps such as quality control, normalization, feature selection, dimensionality reduction, clustering, differential expression (DE) analysis, cell annotation, and trajectory inference. While numerous methods and tools are available for each analysis step, integrating them into a cohesive workflow remains a challenge [2].

Current single-cell analysis tools are available as command-line packages and/or web-based platforms. Command-line packages offer extensive analytical capabilities but require coding and bioinformatics skills [3–6]. Web-based platforms make single-cell analysis more accessible to life scientists by providing an intuitive graphical user interface. These tools can be installed on local computers [7–19], or are available as public servers [20–29]. The analysis capabilities of these platforms vary substantially. Some platforms focus solely on data visualization (e.g., iSEE, ShinyCell, SCope, UCSC Cell Browser) or cluster and DE analysis (Loupe Browser, CellxGene, SPRING, ACT, scQuery, Spectacle). Others offer more comprehensive workflows, from data pre-processing to clustering, DE analysis, and other analytical steps (Cellenics, Asc-Seurat, SCTK2,

ASAP, Cellar, ezSingleCell, NASQAR, ICARUS, SingleCAnalyzer, Shaoxia). Despite the extensive functionalities provided by these platforms, important challenges persist. It is particularly challenging to integrate and analyze large datasets that require coordinated efforts/collaborations and customized analysis workflows to obtain robust results and valid conclusions.

Here we introduce CytoAnalyst, a new web-based platform that offers numerous advantages over existing tools for scRNA-Seq analysis. First, instead of imposing a rigid workflow, the platform enables custom pipeline configuration using an efficient study management system and seven analysis modules (embedding, clustering, DE analysis, gene set management, enrichment, annotation, and trajectory inference). Researchers can navigate between analysis modules and start/resume any analysis steps without losing results or visualization configuration. Second, the platform supports parallel analysis instances, facilitating the comparison of methods or parameter settings available at each step. Third, the advanced sharing system facilitates real-time interaction among team members and seamless analysis continuation across different devices. Access permissions for a shared study can be controlled at granular levels, allowing collaborators to view or alter parameter settings and analysis pipelines. Finally, the grid-layout visualization system supports simultaneous display of different data aspects, allowing for the comparison of multiple labels and plots side-by-side for comprehensive data insights, with the ability to save and reload visualization settings at any analysis step. The platform incorporates multiple blending modes, allowing users to combine plots in various ways for comprehensive data exploration.

To make CytoAnalyst accessible to all researchers, we host it on a high-performance infrastructure with optimized networking and storage capabilities (Dual AMD EPYC 9654 96-Core Processor, 192 cores and 384 threads, 3TB DDR5 RAM, 112TB usable NVMe SSD storage, 4 NVIDIA H100 GPUs). The platform supports all major web browsers without installation or registration requirements.

## 2 Implementation and Methods

### 2.1 Functional Overview

Figure 1 shows the overall structure of CytoAnalyst. The platform consists of three main systems: 1) study management and data sharing, 2) grid-layout visualization system, and 3) core analysis system with seven analytical modules. Here we summarize each system and then describe them in detail in the following sections.

**Figure 1:**
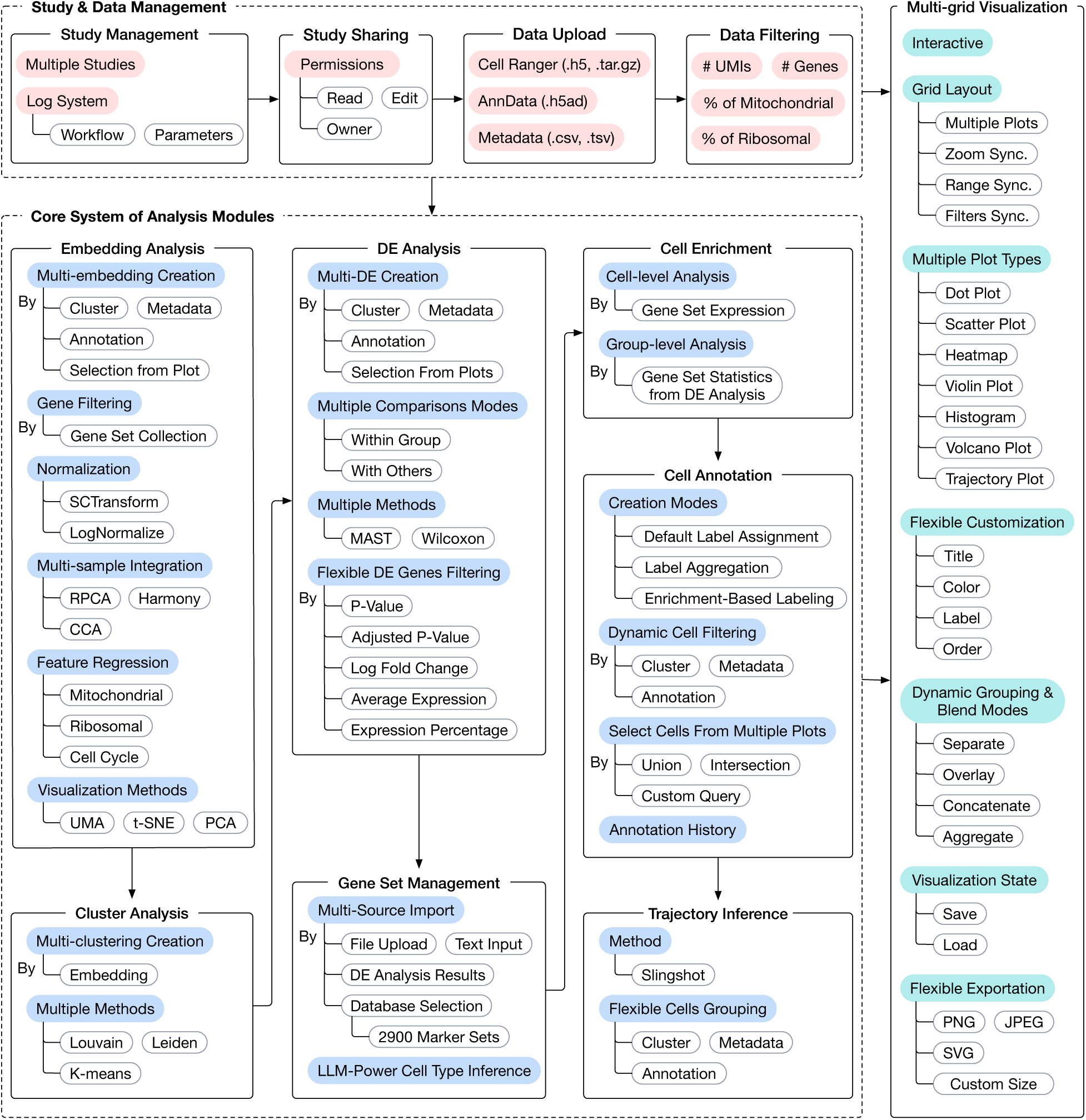
Overview of the CytoAnalyst’s capabilities and core analytical components. The platform consists of three main modules: 1) study management and sharing system, 2) grid-layout visualization system, and 3) core analysis system of seven analytical modules. The management system allows users to create and manage projects and data. The system supports 10X Genomics Cell Ranger in .tar.gz or .h5 format, and Anndata objects in .h5ad format, as well as additional metadata in .csv/.tsv format. Following data upload, users can perform quality control and preprocessing before performing downstream analyses. All analysis parameters and results are automatically logged and can be shared with collaborators through a secure sharing system. The grid-layout visualization system supports flexible and simultaneous display of multiple plot types: scatter plots, violin plots, dot plots, heatmaps, histograms, volcano plots, and trajectory plots. The platform incorporates multiple blending modes, allowing users to combine plots in various ways for comprehensive data exploration. The core system consists of seven analysis modules: embedding analysis, clustering, differential expression (DE) analysis, gene set management, cell enrichment, cell annotation, and pseudo-time trajectory inference. These analysis modules are interconnected, enabling users to seamlessly transition between different steps while maintaining complete control over parameter settings. Advanced users can skip any steps and begin with any process based on their research needs and dataset characteristics.

First, the study management and data sharing system allows users to create and manage their projects. The system supports 10X Genomics Cell Ranger in .tar.gz or .h5 format, and Anndata objects in .h5ad format, as well as additional metadata in .csv/.tsv format. Following data upload, users can perform quality control and preprocessing before downstream analyses. To enhance collaboration and reproducibility, CytoAnalyst incorporates sharing capabilities and maintains detailed analysis logs. Researchers can distribute both data and analysis outcomes with colleagues, while all analysis parameters are automatically documented. The platform provides extensive documentation and tutorials at each analytical step.

Second, the grid-layout visualization system enables dynamic exploration of single-cell data through multiple complementary approaches. The system supports flexible and simultaneous display of multiple plot types: scatter plots, violin plots, dot plots, heatmaps, histograms, volcano plots, and trajectory plots. The platform incorporates multiple blending modes, allowing users to combine plots in various ways for comprehensive data exploration. A distinctive feature of CytoAnalyst lies in its emphasis on interactive visualization and real-time analysis. The platform implements a flexible visualization framework that facilitates the creation of customized plots with adjustable parameters, overlay of multiple data types, and interactive cell selection for focused examination.

Finally, the core analysis system consists of independent modules for embedding analysis, clustering, DE analysis, gene set management, cell enrichment, cell annotation, and pseudo-time trajectory inference. The modules are interconnected, enabling users to seamlessly transition between different steps while maintaining complete control over parameter settings. Advanced users can skip any steps and begin with any process based on their research needs and dataset characteristics. For computationally demanding tasks, CytoAnalyst employs an advanced job queuing system that efficiently manages server resources while delivering real-time progress updates. Users can initiate multiple analyses without waiting for previous tasks to complete. This functionality allows researchers to explore different parameter configurations simultaneously. All analytical results are securely stored on the server and remain accessible through the platform’s interface.

### 2.2 Study Management and Sharing

#### 2.2.1 Study management

The study management system allows users to efficiently manage, share, and monitor their analyses. Researchers can create multiple studies within the platform to organize their data which can be from different experiments or conditions. Each study maintains an independent analysis workflow, allowing focused investigation of related datasets. Users can transition among studies seamlessly through a user-friendly interface, with all analysis results and parameters preserved within their respective contexts.

The platform also provides a secure sharing system for efficient collaborations. Individual studies can be shared through protected links, enabling real-time interaction among team members. Access permissions for a shared study can be controlled at granular levels, allowing collaborators to view or alter parameter settings and the analysis pipeline. The owner of a study can grant or revoke access to the study at any time, ensuring data security and privacy. The sharing system also facilitates seamless analysis continuation across different devices.

To ensure reproducibility and accountability, CytoAnalyst maintains a comprehensive log of all analytical operations. Users can review the analysis history for each study, encompassing data upload details, preprocessing steps, parameter settings, analysis workflow, and visualization. This logging mechanism guarantees that all analytical decisions are documented and retrievable if necessary.

#### 2.2.2 Data upload and pre-processing

CytoAnalyst accepts two standardized formats for single-cell data: output from 10X Genomics Cell Ranger [30] in .tar.gz or .h5 format, and Anndata [31] objects in .h5ad format. Researchers can supplement each dataset with metadata (sample information, experimental conditions, etc.). Multiple files can be uploaded for the same study, enabling integrated analysis across samples or conditions. Upon data selection, CytoAnalyst performs automatic format detection and validation to ensure data consistency. The platform generates an interactive preview of uploaded data, allowing verification of gene identifiers, cell barcodes, and metadata fields.

Following data upload, users can perform quality control and pre-processing before other downstream analyses. The platform computes and visualizes key quality metrics, including unique gene counts per cell, unique molecular identifier (UMI) counts per cell, percentage of mitochondrial genes and/or ribosomal genes. These metrics are displayed as interactive violin plots showing value distributions across all cells. Users can dynamically adjust filtering thresholds while observing effects on cell populations in real time through violin plots and dimensional reduction visualizations.

For multi-sample experiments, quality metrics are computed and displayed independently for each sample, enabling sample-specific quality control thresholds. Sample identity information is preserved throughout the analysis to facilitate downstream data integration and comparative analyses. After initial data processing, users retain the flexibility to incorporate additional samples into the study or supplement existing samples with new metadata. This adaptive approach allows seamless progression to subsequent analysis steps, including data integration, dimensionality reduction, clustering, cell type annotation, differential expression analysis, and trajectory inference.

### 2.3 Grid-layout Visualization

Figure 2 demonstrates the visualization capabilities of CytoAnalyst that enable dynamic exploration of single-cell data through multiple complementary approaches. The visualization architecture centers around a grid-layout system for the simultaneous display of different data aspects (Figure 2A1–8). The framework supports diverse visualization types, including scatter plots, violin plots, dot plots, heatmaps, trajectory plots, histograms, and volcano plots. Each plot is optimized for specific analysis contexts and data types.

**Figure 2:**
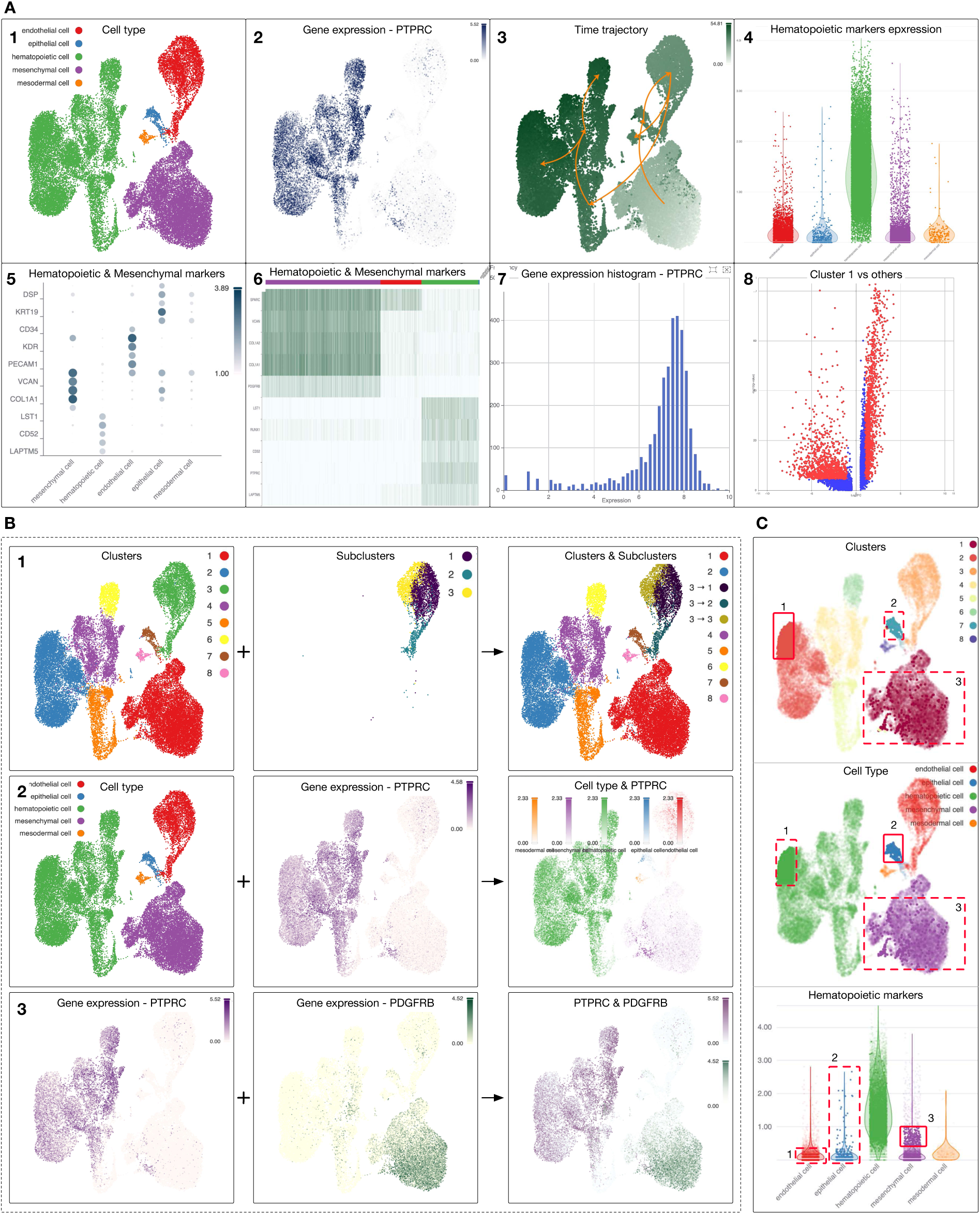
Interactive visualization in CytoAnalyst. A) Chart types supported by the platform, including 1) scatter plot for categorical data (e.g., metadata, clustering, annotation labels), 2) scatter plot for continuous data (e.g., expression, enrichment scores), 3) graph plot overlaying scatter plot, 4) violin plot, 5) dot plot, 6) heatmap, 7) histogram, and 8) volcano plot. B) Blending modes in CytoAnalyst, including 1) aggregate mode for combining two categorical variables, 2) aggregate mode for combining a categorical and a continuous variable resulting in a new plot with a gradient color mapping for each category, and 3) overlay mode for transparent overlaying of two plots for comparison. C) Cell selection across multiple plots using the grid layout, enabling focused examination of specific cell populations. In this example, there are 3 sets of cells selected in 3 different plots. Red solid-lined boxes indicate the selected area/cells in each plot. Red dashed-lined boxes indicate the corresponding selected area/cells in other plots.

Scatter plots facilitate visualization of both continuous and categorical variables using two-dimensional spaces from t-SNE [32] and UMAP [33]. These variables encompass metadata, gene expression levels, cluster assignment, cell type annotation, enrichment results, and trajectory information (Figure 2A1–2). Trajectory plots visualize inferred cellular paths and associated gene expression dynamics. These plots can overlay gene expression levels, pseudotime ordering, or cluster assignments to provide comprehensive views of biological processes (Figure 2A3). Violin plots represent gene expression distributions across cell populations. Users can examine individual genes or gene sets, with options to group cells by categorical variables or their combinations (Figure 2A4). Dot plots and heatmaps excel at visualizing expression patterns across multiple genes and cell populations. Heatmaps provide detailed expression patterns while dot plots offer concise representations highlighting informative genes with their expression levels and percentages (Figure 2A5–6). Histograms display the distribution of individual variables across all cells. Users can adjust bin sizes and range to focus on specific distribution aspects (Figure 2A7). Volcano plots illustrate differential expression results with adjustable significance thresholds and effect size cutoffs (Figure 2A8).

The platform incorporates multiple blending modes, allowing users to combine plots in various ways for comprehensive data exploration. Figure 2B1–3 shows three examples: 1) two categorical variables are aggregated to create a new plot with hierarchical labels, 2) a categorical and a continuous variable are aggregated to create a new plot with a gradient color mapping for each category, allowing for adjusting filtering on each category independently, and 3) an overlay mode for transparent overlaying of two plots for comparison. Overall, supported blending modes include replace, separate, aggregate, overlay, and concatenate. Replace mode enables straightforward feature visualization by substituting plots in the grid layout. Separate mode facilitates side-by-side comparison through the addition of new plots. Aggregate mode combines multiple variables into unified visualizations. For categorical variables, the system generates plots with combined variable coloring. For continuous variables, it calculates and displays average values. When variable types are mixed, i.e., continuous and categorical, the system separates data points by categories and colors them using a gradient scale to allow users to compare values across categories. Overlay mode enables transparent plot overlaying for examining relationships between variables. Concatenate mode stacks multiple features in heatmaps and dot plots for direct comparison.

All visualizations support interactive features including zooming, panning, cell selection, and data point tooltips. Data filtering can be applied to any categorical or continuous variable, affecting individual plots or the entire grid layout. Figure 2C demonstrates the cell selection across multiple plots using the grid layout in CytoAnalyst. When there are multiple plots in the grid layout, users can select cells in one plot and see the corresponding selection in other plots. In this example, there are three different plots with a selected region indicated by a red solid-lined box in each plot. Two red dashed-lined boxes in each plot indicate the corresponding selected cells in other plots. This feature is particularly useful for examining feature expression patterns across multiple visualizations or selecting cells for annotation using complex criteria, such as union or intersection of multiple regions.

Plot arrangements can be modified through drag-and-drop interactions with real-time updates. Users can synchronize zoom levels across plots for direct feature comparison and modify individual plot parameters without altering source data. Color mapping in CytoAnalyst supports both categorical and continuous variables through predefined color presets and custom color palettes and gradients. For categorical variables like cluster assignments, users can define custom colors for each category. For continuous variables, adjustable color gradients for min and max values are available. Users can export any figure in PNG, JPEG, or SVG formats with user-defined dimensions and resolution. They can save and load visualization settings as profiles for future use or collaboration with all parameters and configurations preserved. Users can load saved visualization profiles with only a few clicks.

### 2.4 Core Analysis Modules

At the core of its capabilities, CytoAnalyst consists of seven analysis modules: embedding analysis, clustering, DE analysis, gene set management, cell enrichment, cell annotation, and pseudo-time trajectory inference. These modules are interconnected, allowing for flexible analyses and collaborations. Depending on data and research goals, advanced users can start the analysis using any module, and subsequently refine the results based on their preferred analysis pipelines.

#### 2.4.1 Embedding Analysis

Figure 3 shows CytoAnalyst’s embedding analysis workflow for one or multiple samples. The workflow is compatible with that of Seurat for data integration and dimensionality reduction. Users start the embedding analysis by selecting the cells of interest using sample information, metadata variables, cluster labels, existing cell annotation, and manual selection. Users can filter cells based on any categorical or continuous variable, or directly select the cells from the visualization interface. In parallel, the gene filtering option offers additional analytical flexibility by allowing users to focus their analyses on pre-defined gene sets. This is particularly useful when focusing on specific cell types using known marker genes.

**Figure 3:**
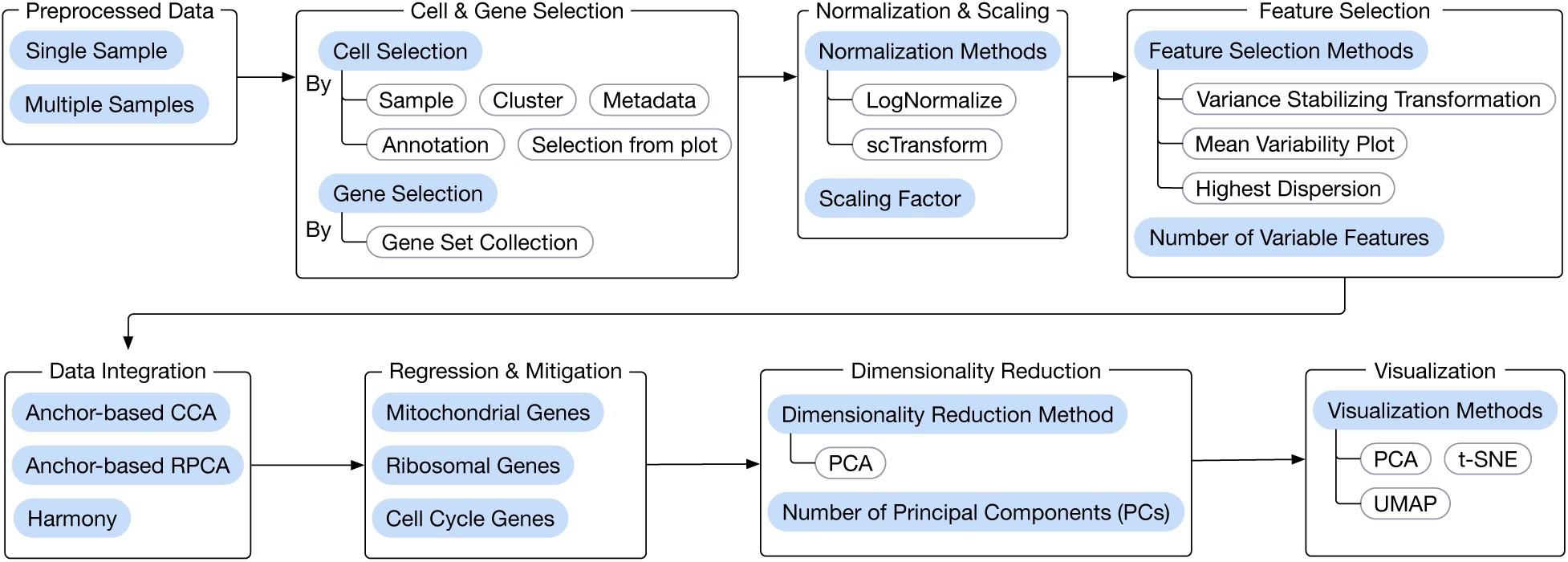
Embedding analysis for one or multiple samples. Users start by selecting the cells of interest using sample information, metadata, cluster labels, existing annotations, and different visualization plots. They can also choose to focus on a pre-defined set of genes using custom gene sets or cell type markers. After cell and gene selection, the next step is to scale/normalize the data (using LogNormalize or scTransform) and to identify highly variable features/genes (using variance stabilizing transformation, mean variability plot, or highest dispersion). The platform includes three integration methods (anchor-based CCA, anchor-based RPCA, and Harmony) for integration and batch correction of single-cell data and samples obtained from different experiments or sources. Users can also regress out unwanted sources of biological variation related to mitochondrial, ribosomal, or cell cycle genes. Finally, users can perform dimensionality reduction and transcriptome landscape visualization using PCA, t-SNE, or UMAP. The platform also facilitates the rapid creation of multiple embedding analyses for categorical variables, such as generating separate embeddings for each cluster. This capability proves particularly valuable for sub-clustering analysis, as global embeddings may not capture the hierarchical data structure.

After selecting cells and genes of interest, users can normalize the data using LogNormalize [34] or SCTransform [35]. They can perform feature selection to identify highly variable genes using Variance Stabilizing Transformation [36], Mean Variability Plot [37], or Highest Dispersion [38]. When analyzing samples and data from multiple sources, users can perform data integration and batch correction using three established approaches named Anchor-based CCA Integration [34], Anchor-based RPCA Integration [39], or Harmony [40]. Users can also regress out unwanted sources of biological variation related to mitochondrial, ribosomal, or cell cycle genes. Finally, users can perform dimensionality reduction and visualize the embedding results using PCA [41], t-SNE [32], or UMAP [33]. The interactive interfaces enable customization of advanced parameters, allowing users to maintain complete control over the analysis.

To extend embedding functionalities, the platform also facilitates the rapid creation of multiple embedding analyses for categorical variables, such as generating separate embeddings for each cluster. This capability proves particularly valuable for sub-clustering analysis, as global embeddings may not capture the hierarchical data structure. With the built-in grid-layout visualization ability, generated embeddings can be visualized with any existing labels. These embeddings serve as foundations for downstream analyses, including clustering, cell type annotation, and trajectory inference.

#### 2.4.2 Cluster analysis

CytoAnalyst implements a flexible cluster analysis framework with multiple algorithms to identify distinct cell populations. The platform includes Louvain [42], Leiden [43], and K-means [44] as clustering methods to accommodate diverse data structures. Louvain and Leiden utilize adjustable resolution parameters to control cluster granularity, while K-means requires the specification of cluster numbers. The graph-based Louvain and Leiden methods excel at global-level clustering analysis for identifying cellular populations across complete datasets, while K-means is more suitable for sub-clustering where the number of clusters is known [45].

To perform clustering, users can choose one or multiple embeddings from embedding analysis and specify the clustering algorithms with corresponding parameters. The platform enables the simultaneous creation of multiple clustering analyses, facilitating the comparison of different parameter settings and algorithms. This capability proves essential for identifying optimal parameters and cluster numbers. The grid-layout visualization system enables direct comparison of clustering results across different embeddings and parameters.

For investigating finer population structures, CytoAnalyst supports hierarchical sub-clustering analysis. Users can perform embedding analysis on specific clusters from initial clustering results, followed by subsequent clustering analysis on these focused embeddings. This approach reveals heterogeneity within major cell types that may be obscured in global embedding spaces.

Once the cluster analysis is done, the cluster labels can be transferred to other embeddings for visualization. In other words, one can use specific embedding for clustering and then choose any other embeddings to display the cell labels. This flexibility enables visualization of sub-clustering results within global embedding spaces alongside primary cluster labels. The platform’s aggregation blend mode facilitates the visualization of multiple clustering levels in unified plots, providing comprehensive views of cellular hierarchies. This visualization approach extends to combining clustering results with metadata labels, enabling the exploration of relationships between cell populations and experimental conditions.

#### 2.4.3 Differential expression (DE) analysis

Figure 4 shows an example DE analysis using the user interface implemented in CytoAnalyst. Figure 4A shows an example DE analysis configuration in which users can choose to compare cells from different clusters (by cluster), cell groups separated by conditions or other variables (by metadata), annotated cell types (by annotation), or customized groups (custom). Via comparison mode, users can choose to compare each cell group against all other groups (with others), or compare cells from different conditions (within cluster). The cell filtering setting allows users to refine the comparative analysis by choosing samples, metadata, cluster labels, and annotated cell types in each of the two groups involved. The method configuration setting allows users to choose the method (Wilcoxon, MAST) and other important parameters (max cell, min percent, log fold-change). Figure 4B shows the preview table with details of the cell groups involved (cell count, selected samples, and clusters), and the total number of cells. Figure 4C shows DE analysis results for a comparison, in which the results can be sorted and/or filtered using any of the computed statistics of the genes (p-value, log fold-change, average expression, percentage of cells with positive expression, and percentage difference). Figure 4D displays the volcano plots in which genes with adjusted p-values less than 5% and absolute log fold-change higher than 2 are highlighted in red.

**Figure 4:**
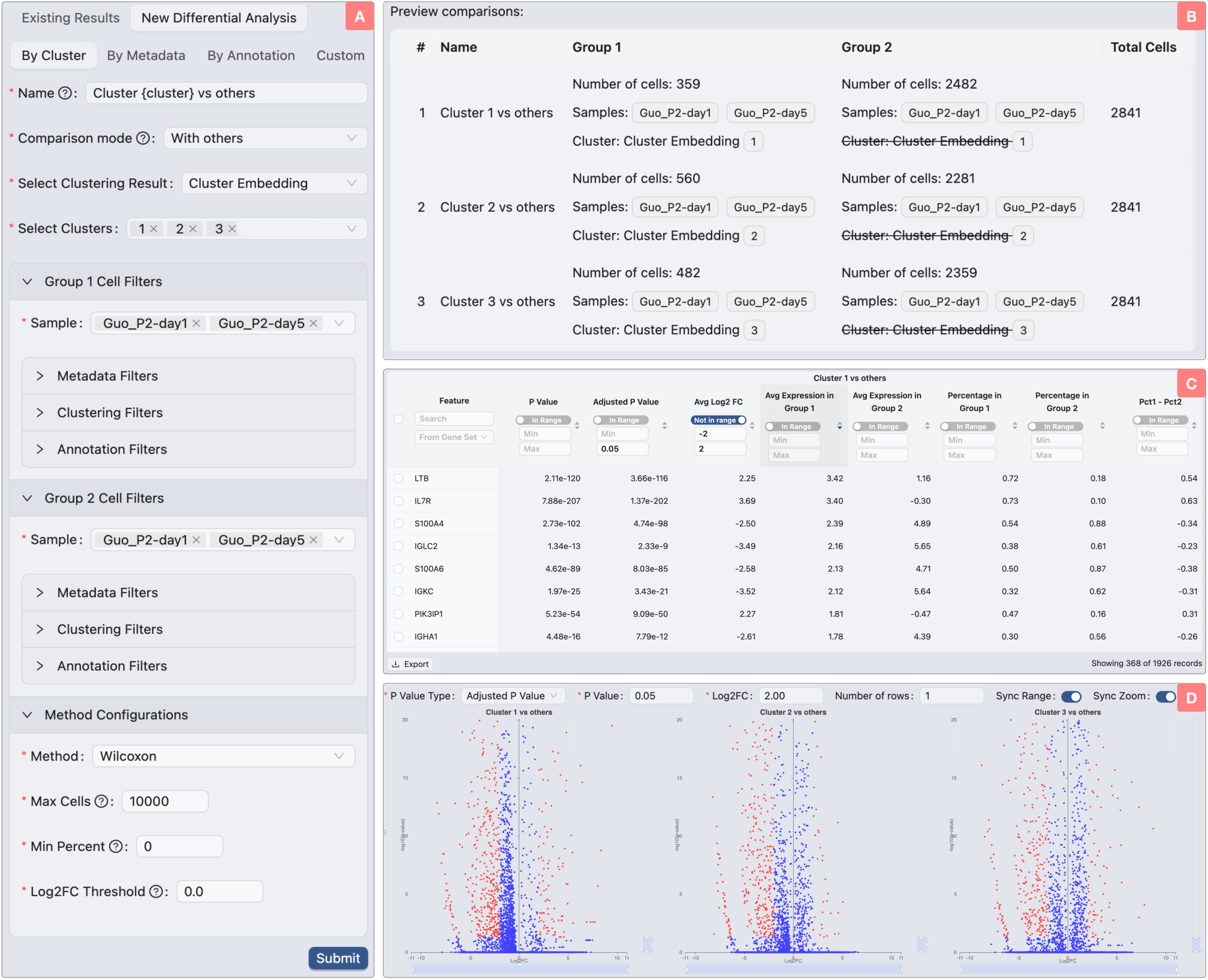
User interface for DE analysis. A) DE analysis configuration. Users can choose to compare cells from different clusters (by cluster), cell groups separated by conditions or other variables (by metadata), annotated cell types (by annotation), or groups manually selected by users (custom). In this example, we perform DE analysis by comparing cell groups obtained from cluster analysis. Via comparison mode, users can choose to compare each cluster against all other clusters (with others), or compare cells from different conditions (within cluster). The cell filtering setting allows users to choose specific samples, metadata, cluster labels, and annotated cell types. The method configuration setting allows users to choose the hypothesis testing method (Wilcoxon, MAST) and important parameters: 1) total number of cells involved (max cell), 2) minimum expression percentage for genes (min percent), and 3) minimum log fold-change for genes. B) Preview table before performing DE analysis. In this example, users are comparing each of the three clusters against all other clusters. The table displays the name for each comparison, details of the cell groups involved (cell count, selected samples, and clusters), and the total number of cells. C) DE analysis results for one of the three comparisons. The results can be sorted and/or filtered using any of the computed statistics of the genes: 1) p-value and adjusted p-value, 2) log fold-change, 3) average expression in each group, 4) percentage of cells in which the genes are expressed, and 5) percentage difference. D) Volcano plots of all comparisons that show the adjusted p-values and log fold-change. In this example, genes with adjusted p-values less than 5% and absolute log fold-change higher than 2 are highlighted in red while the rest of the genes are highlighted in blue.

Overall, we implement a comprehensive framework for comparative analysis among different types/clusters, conditions, and time points. Users can perform DE analysis on any cell subset using an advanced selection system that combines interactive visualization and filtering based on both categorical and continuous variables. The DE analysis workflow supports three distinct comparison modes: between groups, within groups, and user-defined group comparisons. Between-group analysis identifies genes differentially expressed between clusters or categorical variables, such as comparing a cluster against all other cells. Within-group analysis examines condition-specific effects in defined populations, enabling comparison of identical clusters across different conditions. Custom analysis refers to customized grouping defined by users. The platform facilitates the creation of multiple DE analyses simultaneously, providing a preview interface that displays comparison groups with their respective cell counts and gene numbers.

For hypothesis testing, users can choose between Wilcoxon rank-sum test [46] and Model-based Analysis of Single-cell Transcriptomics (MAST) [47] to compute the p-values for the genes, followed by a correction for false discovery rate using Benjamini-Hochberg [48]. After hypothesis testing, users can refine the list of DE genes using p-value, log fold-change, average expression, and minimum expressing cell percentage (in each group or between groups). Result exploration for DE analysis utilizes interactive volcano plots arranged in grid layouts, accompanied by comprehensive statistical tables. The interface enables gene searching, filtering, and highlighting across plots. Selected genes can also be incorporated into existing gene set collection for other downstream analyses.

#### 2.4.4 Gene set management

A gene set is a collection of unique genes grouped together based on a shared characteristic or function. A gene set can represent a cellular process or functional module, signaling pathway, markers of a specific cell type, markers of a condition or disease, or simply a set of DE genes obtained from a comparative analysis. There are multiple applications of gene sets in CytoAnalyst, including: 1) gene filtering in embedding analysis, 2) cell enrichment and annotation, 3) cell type inference using Large Language Models (LLMs), and 4) visualization and identifying patterns of gene expression associated with specific cellular processes or phenotypes.

CytoAnalyst implements a hierarchical system for creating, importing, and managing gene sets. The platform organizes gene sets into collections, enabling logical grouping of closely related gene sets. This hierarchical structure facilitates the management of different biological contexts, such as varying cell type granularity levels or pathway-specific gene groups. Users can create distinct collections for broad cell types, cell subtypes, or biological pathways to explore enriched processes in specific cell populations. The platform supports multiple methods for gene set creation and management. Users can create new collections, import gene sets from external sources in tabular or GMT formats, or choose from curated pre-defined collections. Within each collection, users maintain full control over gene set composition, including renaming, deletion, and modifying the gene set (adding and removing genes).

By default, CytoAnalyst embeds a comprehensive collection of cell type markers available in CellMarker 2.0 [49]. The database contains a curated compilation of experimentally validated markers for known cell types in human and mouse tissues. For humans, users can select from more than 400 tissues, 1,500 cell types, and 15,000 markers. Similarly, the mouse reference collection offers users approximately 300 tissues, 1,400 cell types, and 12,000 markers. Users can easily search for these reference gene sets (by tissue or cell type) and incorporate them into their analysis pipelines with minimal effort.

The gene set management interface also provides a cell-type inference tool that leverages LLMs to infer potential cell types associated with a gene set. Given the tissue and the gene set (marker genes), the tool returns a list of cell types most likely present in the sample, structured in a cell ontology hierarchy. The inference tool is based on Meta’s llama 3.3 [50], a recent 70B parameter model, using the Ollama framework to create a responsive API system that enables efficient LLM communication. To ensure an accurate inference with a consistent output format, we design a prompt template that directs the LLM with specific guidance and context.

#### 2.4.5 Cell enrichment analysis

CytoAnalyst supports both cell-level and group-level enrichment analyses. Cell-level enrichment basically performs enrichment analysis of pre-defined gene sets for each individual cell. Given a cell, CytoAnalyst first calculates the z-score of each gene and then compares the z-scores of genes in a gene set against genes in all other gene sets using one-sided Welch’s t-test [51]. In addition to the p-value for each gene set, the software also returns a score (average z-score of genes in the gene set), and score difference (by subtracting the average z-score of genes in the gene set from the average z-scores of genes in other gene sets). Users can visualize the p-values, scores, score differences across cells and gene sets for cell annotation or developmental stages.

In addition to cell-level enrichment, CytoAnalyst also supports enrichment analysis for a group of cells (group-level enrichment). Using the DE analysis results, the platform computes the p-values and statistics of pre-defined gene sets for the cell group. The platform first ranks the genes using log2FC and p-values and then applies FGSEA [52] to compute enrichment scores and statistical significance for each gene set. The platform also calculates the enrichment score difference as the difference between the enrichment score of the target gene set and the average enrichment score of all other gene sets. The obtained enrichment score, enrichment score difference, and statistical significance are then assigned to all cells of the underlying group.

Cell enrichment primarily facilitates cell type annotation and cell state identification. For example, users can enrich cells with known cell type markers to assign labels to distinct populations, or enrich DE genes with biological pathways to characterize cellular states. In both cases, the platform calculates enrichment scores for the target gene set, enrichment score differences between the target gene set and other genes, and the statistical significance for individual cells or cell groups.

#### 2.4.6 Cell annotation

Cell annotation is a critical step in scRNA-Seq analysis, enabling researchers to assign meaningful labels to distinct cell populations. CytoAnalyst implements a flexible annotation system combining manual curation, marker-based enrichment, and LLM-based inference for cell type assignment. Users can create a new annotation instance through three available options: 1) default label assignment (Figure 5A), 2) label assignment using metadata, clustering, and former annotations (Figure 5B), and 3) assignment using cell enrichment results (Figure 5C). In the first option (default), users can assign an unknown label to all cells in the dataset and then gradually refine the annotation through manual examination and analysis. In the second option (label aggregation), users can combine variables from metadata, clustering, and former annotations to generate new cell labels. In the third option (enrichment analysis), users can assign initial cell labels based on enrichment statistics, metadata, clustering, and other variables.

**Figure 5:**
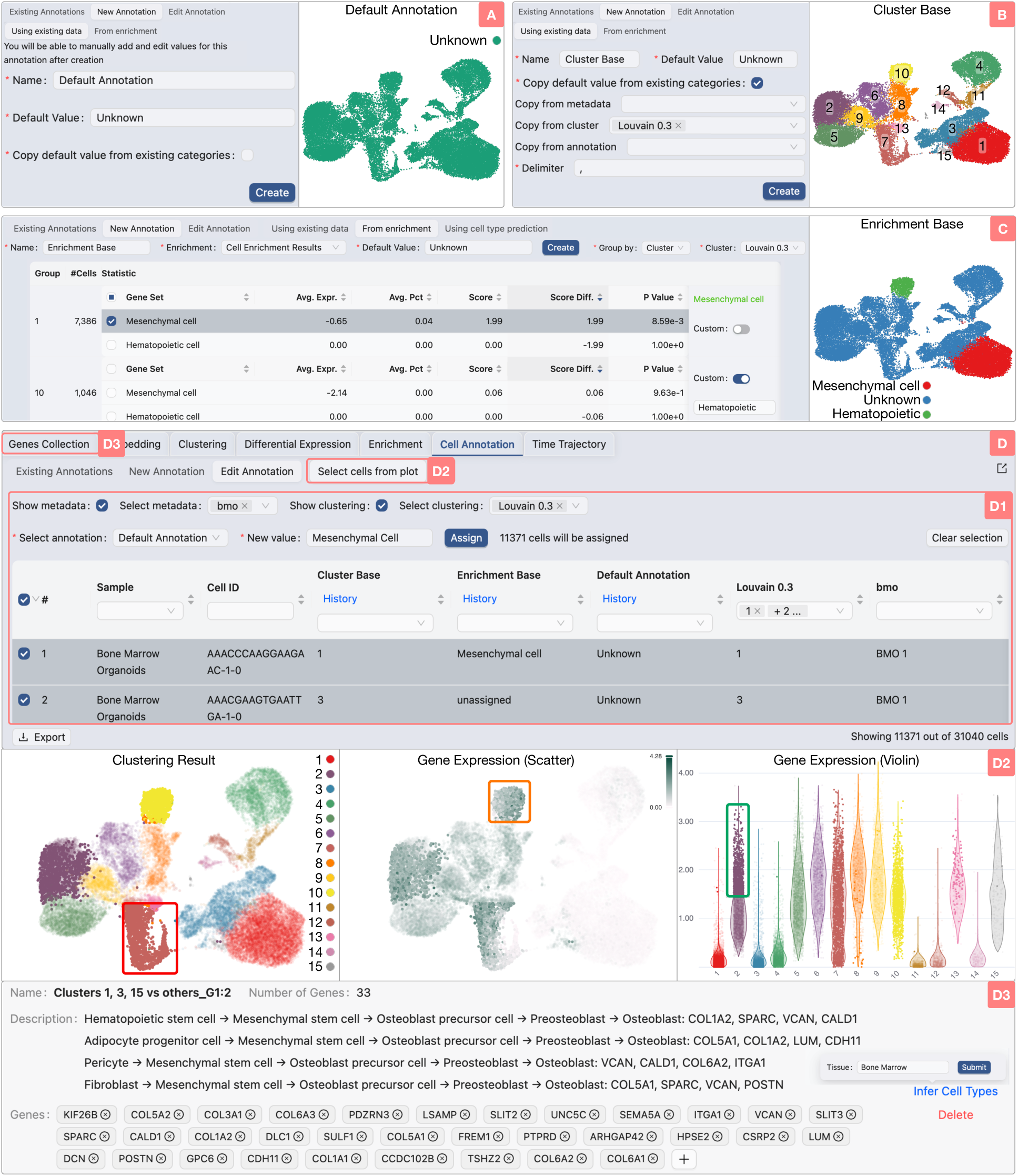
Cell annotation interface. A) Default annotation initialization. Users can create a default annotation in which all cells are unknown, and then gradually refine the annotation. B) Annotation initialization using clustering results, metadata, and/or former annotation. C) Annotation initialization using enrichment analysis results. Using enrichment results, users can assign labels to a group of cells by either selecting a gene set with satisfying statistics in the group or assigning a custom label to cells in a group. D) Annotation editing interface. CytoAnalyst allows users to flexibly choose and assign new labels to cells (D1). They can view all cell labels in the annotation table. In addition, users can choose to display specific information from metadata (show metadata) or clustering results (show clustering) as columns. In this example, we select 11,371 cells (out of 31,040) from clusters 1, 3, and 15, and then assign them as Mesenchymal (new value). Moreover, users can select the cells from different plots (D2). They can also use the LLM tool to infer the cell type based on the markers of the cells (D3).

The annotation feature is not an isolated module but supplemented by other modules, including *Gene Set Management* and *Cell Enrichment Analysis* so that users can create an accurate and informative annotation (see sections above). Users can switch between the annotation interface and the interface while editing the annotation. Figure 5D shows the interface for annotation editing. The interface allows users to flexibly filter cells using metadata, clustering, or any other labels (Figure 5D1). The platform displays cells in a tabular format with current annotations, metadata variables, and cluster information. This table supports additional filtering operations and cell export capabilities. The annotation interface also integrates with the grid-layout visualization system, allowing users to select cells from multiple plots (Figure 5D2). For complex selections across multiple plots, users can define how to combine these regions using union, intersection, or custom queries that support AND, OR, and NOT operators. CytoAnalyst also provides users with a way to infer potential cell types for each group of cells based on their marker genes and tissue information (Figure 5D3). By switching to the Genes Collection interface, users can utilize an LLM model to infer the cell type based on the marker genes of a cell group. Finally, users can assign new labels to the selected cells, and the platform will update the annotation results immediately. For each session, the platform maintains the annotation history, allowing users to review and revert previous assignments.

#### 2.4.7 Trajectory inference

Trajectory inference in single-cell analysis is a computational approach that models dynamic biological processes, such as cellular differentiation, developmental pathways, or disease mechanisms, by mapping the sequential gene expression changes cells undergo temporally [5]. Cytoscape utilizes Slingshot to infer cell lineage trajectories and pseudo-temporal ordering from single-cell RNA-seq data [53]. The method has been shown to perform robustly across diverse datasets, balancing accuracy and flexibility in identifying branching lineages [54]. Slingshot treats cell clusters as nodes in a graph and constructs a minimum spanning tree (MST) connecting these nodes, thereby identifying the global lineage structure (e.g., the number of lineages and branching points). Next, it assigns pseudo-times to individual cells by fitting principal curves [55] to the data along each lineage, starting from a user-specified or algorithm-determined root cluster. These curves model smooth trajectories through the cells’ expression space, capturing their progression along divergent differentiation paths.

CytoAnalyst enhances trajectory inference by letting researchers group cells in multiple ways, using clusters, annotated cell types, or metadata (like treatment groups or patient cohorts). This flexibility allows researchers to focus on specific populations, such as tracking differentiation in cells from a particular experiment or comparing trajectories between healthy and diseased samples. Additionally, CytoAnalyst offers customizable settings for Slingshot, including the ability to choose how cluster distances are measured, modify the convergence threshold to control the precision of principal curve fitting, and so on. These customizations help researchers reproduce or apply them to their specific case studies.

After inferring trajectories, CytoAnalyst provides an intuitive visualization for users to explore and interpret the results. Researchers can visualize trajectories and gene expression in overlay mode to observe how a marker gene peaks at a branching point and in aggregate mode to visualize multiple lineages on a single plot or in separate mode to examine individual lineages in detail.

## 3 Implementation

Figure 6 shows the overall architecture of CytoAnalyst. The web-based platform utilizes modern web technologies to provide a seamless user experience across different devices and browsers. At the interface, we use the React framework to enable the creation of dynamic, responsive components that are updated in real time as users interact with the platform. To enable real-time collaboration features (analysis sharing, result updating, etc.) we build the back end using WebSocket along with Meteor, a full-stack JavaScript platform that simplifies the development of real-time web applications. It provides a scalable, reactive architecture that enables efficient data synchronization between clients and servers. With such back-end architecture, CytoAnalyst provides real-time updates during data processing and synchronizes both individual and shared analyses across all clients, sessions, and devices. This enables users to monitor and alter analysis progress, intermediate results, and visualization. In addition, data management is one of the key components of CytoAnalyst’s architecture, CytoAnalyst leverages MongoDB, a scalable NoSQL database for efficient storage and retrieval of large datasets. This robust database infrastructure supports CytoAnalyst’s data management system, including automatic archiving of unused projects to optimize server resources.

**Figure 6:**
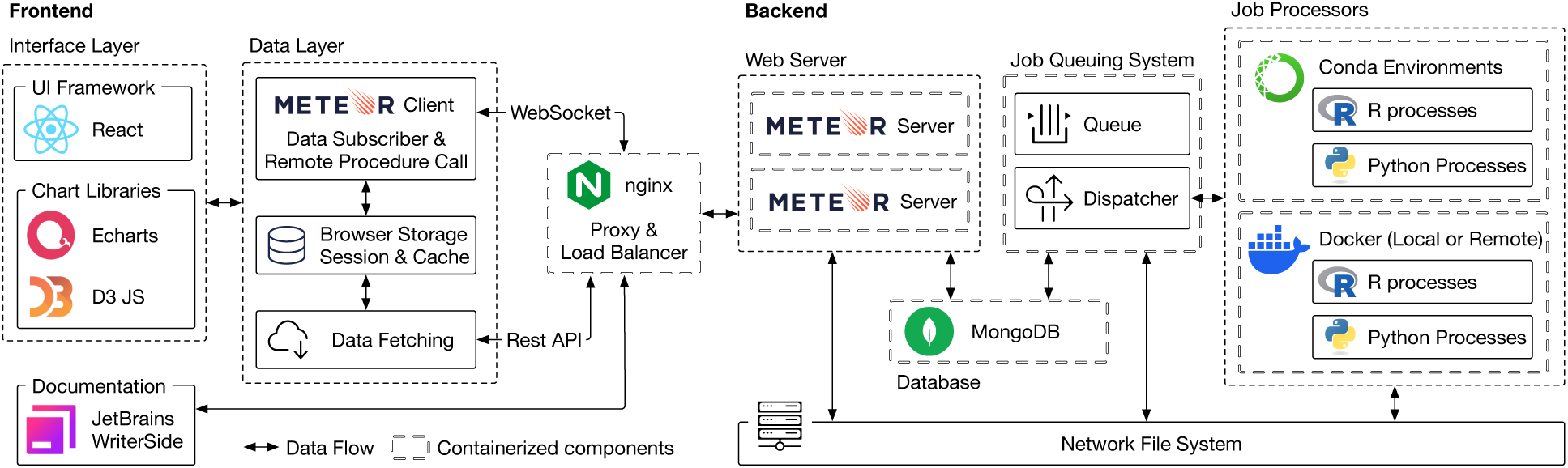
CytoAnalyst software architecture. The platform is built on a modern web stack with React for the frontend interface, Echarts and D3 for interactive visualization, JetBrains WriterSide for documentation, Meteor for the back-end server, MongoDB for database management, and R and Python for data processing and analysis. CytoAnalyst implements a high-performance job queuing system to manage computational tasks and resource allocation, synchronizing real-time progress updates across sessions and devices. The platform is deployed using Docker containerized technologies to ensure portability and scalability, and is hosted on an enterprise-grade server infrastructure with optimized networking, storage, and computational capabilities to handle large-scale single-cell datasets.

The core sharing system in CytoAnalyst utilizes a flexible permission management framework. Project owners can share analyses as read-only, read-write, or full ownership. This granular permission system enables team leaders to maintain appropriate control while facilitating collaborative analysis. For each shared project, CytoAnalyst maintains complete version control. All analytical steps, including parameter settings and computations, are automatically logged and preserved. We implement a token-based authentication system tied to each study, ensuring only authorized users can view or modify analysis results. The platform maintains detailed access logs and enables project owners to revoke sharing permissions when needed. All analytical steps, including parameter settings and computations, are automatically logged and preserved.

CytoAnalyst implements its data processing and analysis modules in R and Python, leveraging popular libraries for single-cell analysis. A custom job queuing system manages the platform’s analytical workflows, providing users with real-time progress updates while efficiently handling computationally intensive tasks like data integration, clustering, and DE analysis without sacrificing performance. The platform builds its job queuing system on a distributed task scheduler that manages multiple concurrent jobs and allocates computational resources dynamically. The task dispatcher inside the job queuing system works with conda environments and docker containers to ensure the scalability and reproducibility of the analysis workflow. Users can create multiple analysis jobs simultaneously and monitor the progress of their jobs in real time through the platform’s interactive interface.

We deploy CytoAnalyst using containerized technologies from Docker to ensure portability and scalability. Docker containers encapsulate each component, enabling seamless deployment across different environments and cloud providers. The containerized architecture allows CytoAnalyst to scale computational resources easily to handle large datasets and high user traffic. To support CytoAnalyst’s architecture, we host it on a high-performance infrastructure with optimized networking and storage capabilities.

## 4 Case studies

As case studies, we analyze three single-cell datasets obtained from previous studies [56–60]. The first dataset consists of 31,040 cells in bone marrow organoids [56]. The second dataset comprises 15,457 cells collected from the sun-protected inguinoiliac region of whole-skin samples from five male donors [57]. The third dataset consists of 5,828 cells from bone marrow that were collected from three experimental studies [58–60].

### 4.1 Case Study 1: Annotation of Bone Marrow Organoids

Here we analyze the single-cell dataset (31,040 cells) from Frenz-Wiessner et al. [56], in which bone marrow organoids were generated from human induced pluripotent stem cells. Through manual examination of both scRNA-Seq and matched flow cytometry data, the authors identified five major cell types. Assuming that we do not know the number or cell types, nor the true annotation of the single-cell data. We aim to identify potential cell types based on the analysis of the single-cell data alone, without using external data from the matched flow cytometry. We will do so following a logical order of analysis steps: transcriptome landscape visualization, cluster analysis, DE analysis, and cell type annotation.

#### Transcriptome Landscape Visualization and Louvain Clustering

To initiate the analysis, we upload the file Bone Marrow Organoids.h5ad obtained from Frenz-Wiessner et al. [56]. Note that the authors have already filtered barcodes with fewer than 400 detected genes, more than 40,000 counts in total, or mitochondrial genes that exceeded 10% of the total number of gene counts. They also employed Scrublet v.0.2.3 [61] with default parameters, filtering transcriptomic profiles with a predicted double score exceeding 0.2 for the removal of doublets.

Figure 7A shows the transcriptome landscape of the updated data using UMAP. The visualization clearly shows that the transcriptome landscape consists of three major cell populations (marked as I, II, and III). To determine potential cell groups, we perform Louvain clustering [42] with the default resolution of 0.3 (Figure 7B). Acknowledging that Louvain clustering has the potential to produce a large number of clusters, we execute the algorithm with different resolutions to explore cluster granularity and to identify robust boundaries (Figure 7C).

**Figure 7:**
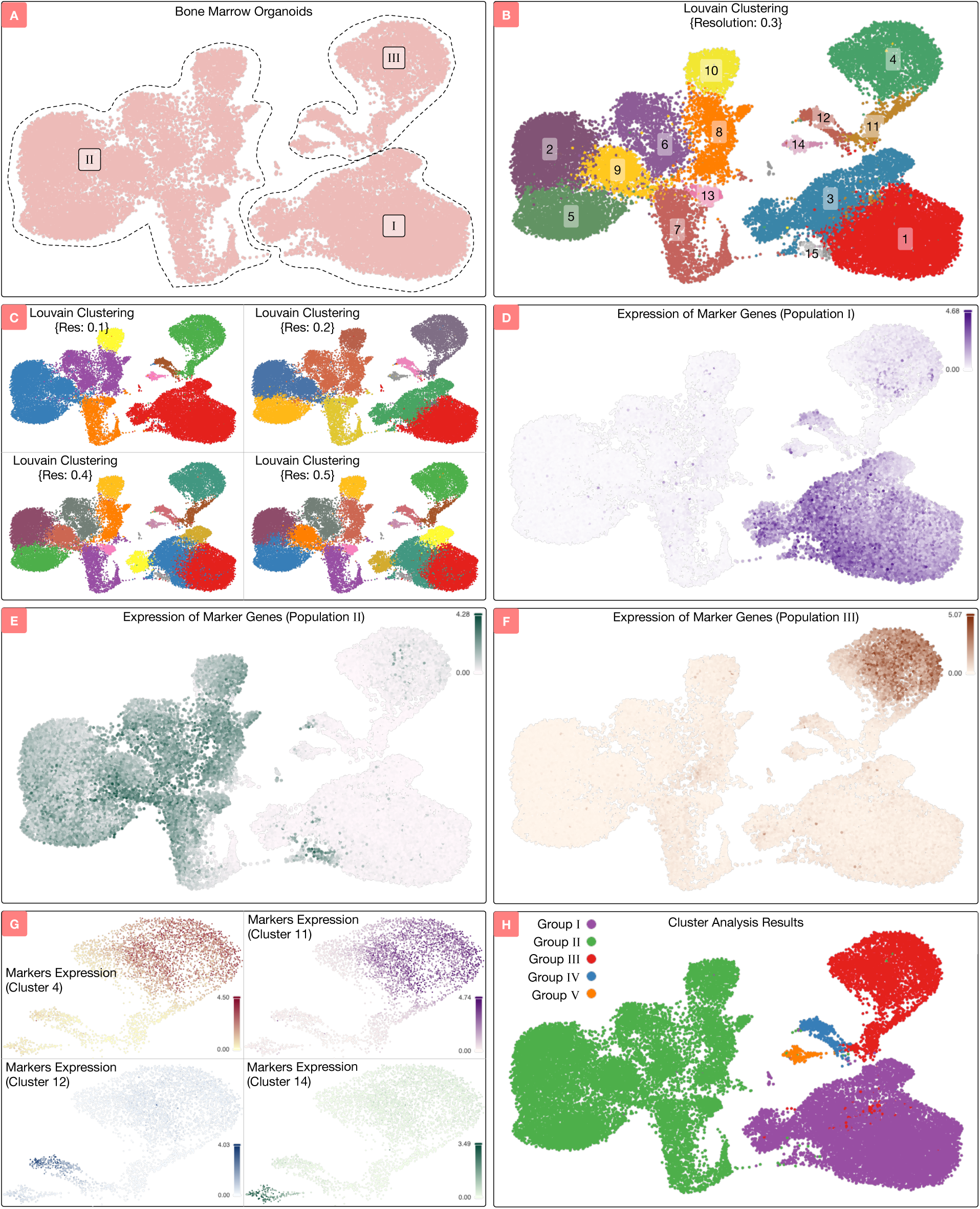
Visualization, Louvain clustering, and differential expression (DE) analysis. A) Transcriptome landscape visualization using UMAP. The landscape can be separated into three populations (I, II, and III) that are highlighted by the three dashed lines. B) Louvain clustering using the default resolution of 0.3. C) Louvain clustering results using resolutions of 0.1, 0.2, 0.4, and 0.5. The next step is to perform DE analysis and visualize the expression of the marker genes across the transcriptome landscape. D—F) Expression of marker genes for populations I, II, and III, respectively. G) Expression of marker genes for clusters 4, 11, 12, and 14 of population III. Final cell grouping based on visualization, Louvain clustering, and DE analysis.

Figure 7B and C show that regardless of resolution settings, Louvain is able to separate the cells into three major populations which is consistent with what we observe from the transcriptome landscape. Louvain with a resolution of 0.3 identifies 15 clusters, in which clusters 1, 3, and 15 correspond to population I; clusters 2, 5, 6, 7, 8, 9, 10, and 13 correspond to population II; clusters 4, 11, 12, and 14 correspond to population III. Similarly, Louvain with resolution 0.1 (panel C) identifies 8 clusters, in which cluster 1 corresponds to population I; clusters 2, 4, 5, and 6 correspond to population II; clusters 3, 7, and 8 correspond to population II. In summary, Louvain analysis results with different resolutions all separate the transcriptome landscape into three major cell populations as we observed in Figure 7A–C. For the next step of the analysis, we proceed with Louvain results using the default resolution of 0.3 (Figure 7B), but all other resolutions are likely to lead to similar annotation results as we will show in the following text.

#### DE Analysis and Cluster Verification

We proceed with the Louvain clustering results using the default resolution of 0.3 (Figure 7B). There are three major cell populations (I, II, and III). Next, we perform DE analysis to identify the marker genes of each population and visualize their expression. The goal is to verify whether we should further divide each population into smaller cell groups. We conduct three DE analyses: population I (clusters 1, 3, 15) versus others; population II (clusters 2, 5, 6, 7, 8, 9, 10, 13) versus others; population III (clusters 4, 11, 12, 14) versus others. We use the Wilcoxon rank sum test to calculate the p-values of the genes and then adjust the p-values using Benjamini-Hochberg [48]. We then use the following criteria to identify the marker genes of each population: 1) log fold change of 3 or higher, 2) FDR p-value less than 5%, 3) average expression more than 1, and 4) the difference in the percentage of cells expressing the gene between groups is at least 50%. We then visualize the expression of the marker genes.

Figures 7D–F show the average expression of the marker genes for each of the three cell populations. The marker genes of population I have high expression in the population and have negligible expression in any other populations (Figure 7D). Therefore, we are confident that the first population consists of a single cell type. Similarly, the marker genes of population II have high expression in the population and have negligible expression in any other populations (Figure 7E). Therefore, it is most likely that population II consists of a single cell type as well. However, the markers of the third population (clusters 4, 11, 12, and 14) have high expression only in cluster 4 and not in clusters 11, 12, and 14. This suggests that cluster 4 represents a distinct cell type from clusters 11, 12, and 14.

Consequently, we perform additional DE analyses for each cluster in population III. Figure 7G shows the expression of marker genes in each of the four clusters. The top two panels show the expression of marker genes in clusters 4 and 11. Interestingly, the expression of marker genes in these two clusters share very similar patterns. Therefore, we merge these two clusters together. The bottom two panels show the expression of marker genes in clusters 12 and 14, in which marker genes of each cluster are highly expressed in their respective cluster but not in other clusters. At the end, we divide population III into three cell groups: clusters 4 and 11 together, cluster 12, and cluster 14. In summary, through visualization, Louvain clustering, and DE analysis, we identify five cell groups. Figure 7H and Table 1 show the final clusters and their respective markers.

**Table 1:**
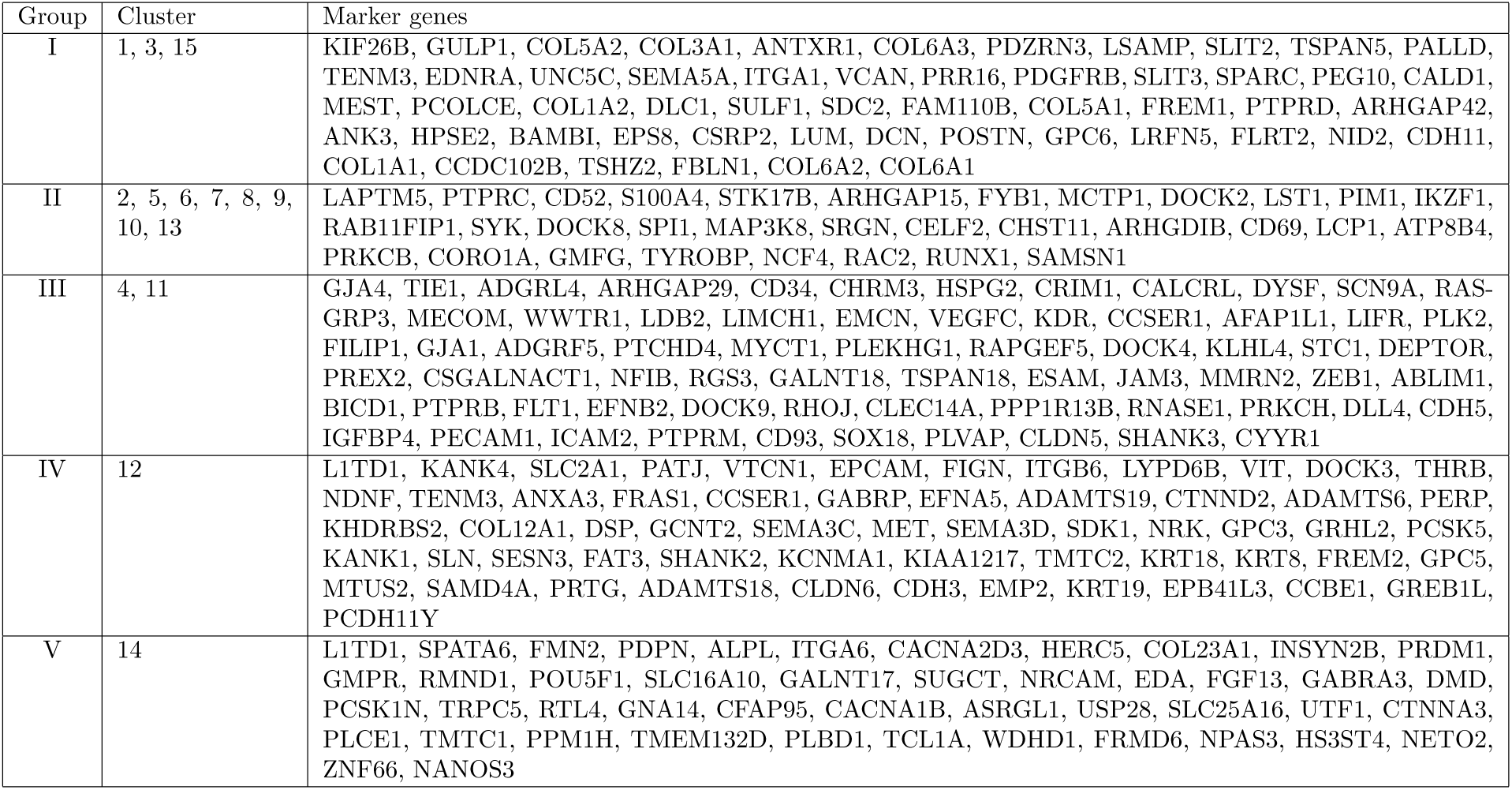
Marker genes obtained from DE analysis. We use the following criteria to identify the genes that are differentially expressed: 1) Log2FC≥2, 2) FDR p-value ≤ 0.05, 3) expression ≥ 1, and 4) percentage of cells expressing the genes between groups ≥ 50% (40% for cluster 12).

#### Cell Type Annotation using built-in LLM-based Inference

Through visualization, clustering, and DE analysis, we identify five groups of cells. Here we aim to assign the cell groups to known cell type labels. For each of the five groups and their associated markers (Figure 7H and Table 1), we use the built-in inference tool to search for potential cell types. Given the tissue and associated markers, the inference tool returns a list of cell types most likely present in the sample, structured in a cell ontology hierarchy. Table 2 displays the top predictions for each cell group, sorted in descending order of likelihood.

**Table 2:**
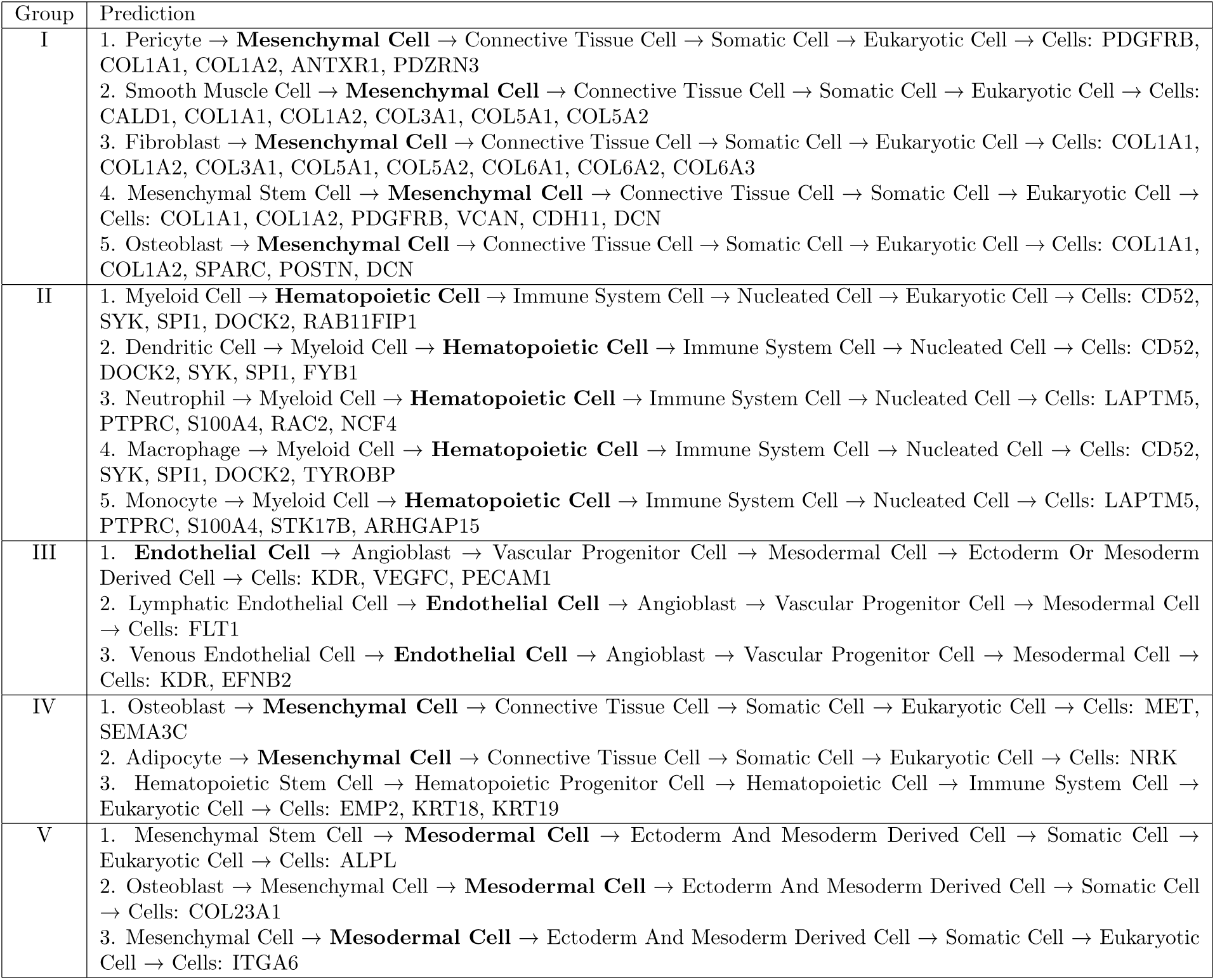
Predicted cell types using the LLM-based inference tool.

Our strategy for cell type assignment is to search for the label that appears most frequently in the top predictions. If there are multiple labels with the same frequency, we choose the cell type with the lowest order (more fine-grained) in the cell ontology hierarchy. The goal is to reduce the assignment error since a more fine-grained label needs more evidence. Based on such strategy, we label Group I as Mesenchymal cells because Mesenchymal appears in most predictions. Similarly, we classify Group II as Hematopoietic stem cells because the top three predictions all point to this cell type. Group III is classified as Endothelial cells as the cell type appears four times in the five predictions. Interestingly, the LLM suggests that Group IV should be assigned to Mesenchymal cells (similar to Group I). Therefore, we merge Group I and Group IV together and label them as Mesenchymal cells. Finally, group V is assigned to Mesodermal cells.

Figure 8 summarizes the complete analysis for this case study. Data visualization shows that there are three major cell populations in the transcriptome landscape (Figure 8A). Following the analysis of Louvain clustering and expression patterns of the marker genes of each population, we determine that the dataset consists of five cell groups (Figure 8B). Using the LLM inference tool, we assign each cell group to a known cell type label with the highest likelihood (Figure 8C). The final annotation determined by CytoAnalyst is highly similar to the annotation provided by the authors (Figure 8D). The two annotations share 99% similarity, with the difference being that CytoAnalyst assigns cells in group IV (blue cells in Figure 8B) to Mesenchymal instead of Epithelial cells. Endothelial cells can undergo a process called endothelial-to-mesenchymal transition (EndMT) where they acquire mesenchymal characteristics. We hypothesize that the authors of the dataset were able to distinguish between the two cell types using external evidence from flow cytometry data [56].

**Figure 8:**
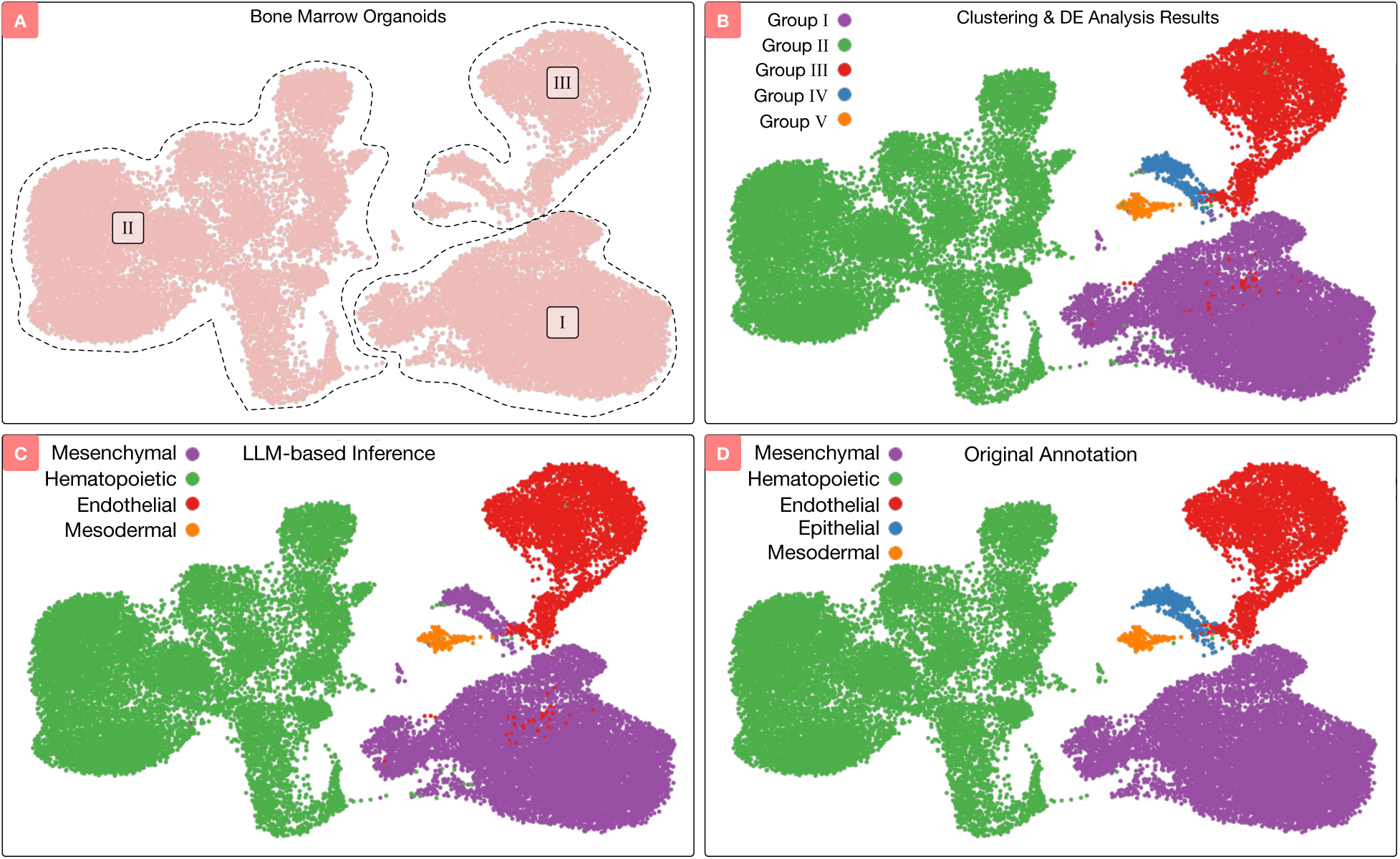
Final cell type annotation results. A) Transcriptome landscape visualization. B) Final grouping using clustering and DE analysis. C) Cell types annotated by CytoAnalyst using the built-in LLM-based inference tool. D) Cell types annotated by Frenz-Wiessner et al. using single-cell and flow cytometry data. The two annotations share high similarity with the only difference is that CytoAnalyst assigns group IV to Mesenchymal instead of Epithelial.

### 4.2 Case Study 2: Cell Type Markers of Skin Samples

We analyze the single-cell data from Soĺe-Boldo et al. [57]. The data has a total of 15,457 cells from whole-skin samples of five male donors: two young donors (25 and 27 years old) and three old donors (53, 69, and 70 years old). The authors performed standard data processing, data integration, and dimension reduction using Seurat [3]. They performed cluster analysis and cell type annotation, resulting in 17 clusters (using Louvain) and 9 main cell types using known cell markers: keratinocytes, fibroblasts, macrophages/dendritic cells, T cells, vascular and lymphatic endothelial cells, pericytes, erythrocytes, and melanocytes. Next, they isolated 5,948 fibroblasts and performed functional enrichment to identify four fibroblast subtypes: secretory-reticular fibroblasts, pro-inflammatory fibroblasts, secretory-papillary fibroblasts, and mesenchymal fibroblasts. The authors also performed DE analysis to identify the markers for the clusters, cell types, and subtypes.

In this case study, we use the DE analysis module from CytoAnalyst we conduct four distinct DE analyses: (1) identification of cluster markers, (2) identification of fibroblast-specific markers in young samples, (3) identification of fibroblast-specific markers in old samples, and (4) comparison of fibroblast subpopulations in young samples. In the first analysis, cluster markers are identified by comparing each cluster against all other clusters. With 17 clusters in total, this results in 17 DE analyses. In the second and third analyses, we compare each fibroblast cluster against all other cells in young and old samples, respectively. With four clusters within the fibroblast population, this results in a total of 8 DE analyses. In the final analysis, we compare each fibroblast subpopulation against other fibroblast subpopulations in young samples, resulting in 4 DE analyses. For all comparisons, we use the Wilcoxon rank-sum test to calculate the p-values.

The parameter settings used for all analyses are shown in Figures S1 and S6. The volcano plots are shown in Supplementary Figures S2 – S5, and S7. The DE genes for each of the four analyses are reported in Supplementary Tables S1–S4, respectively. Overall, the results from CytoAnalyst are consistent with the published results. In the first analysis (17 DE analyses for 17 clusters), most of the markers identified by CytoAnalyst for each cluster are also confirmed by the authors (88.4%). In the second analysis, fibroblast-specific markers identified by CytoAnalyst in young samples share high similarity (91.9%) with the list of markers provided by the authors. In the third analysis, fibroblast-specific markers identified by CytoAnalyst in old samples have 91.1% similarity with the markers identified by the authors. In the fourth analysis, markers identified by CytoAnalyst have 84.7% similarity with the markers identified by the authors. Detailed analysis workflow and results are reported in Supplementary Section 1.

### 4.3 Case Study 3: Trajectory Inference of Bone Marrow Cells

We analyze the bone marrow dataset obtained from Björklund et al. [62] in which the authors integrated the data from three different experiments [58–60]. The authors processed the data using standard Seurat protocol, including cell filtering, normalization, feature selection, dimensionality reduction, and clustering. The authors performed trajectory inference using Slingshot to identify developmental trajectories of bone marrow cells. They also used tradeSeq [63] to identify genes that change during the developmental trajectories of each lineage: Cd34 (Hematopoietic stem/progenitor cells, or HSPC), Ms4a1 (B cells), Ltf (Granulocyte cells), and Siglech (Dendritic cells). The authors published the lineages obtained for only HSPC, which can serve as a reference for our analysis. Here we aim to reproduce the reported HSPC lineages using the same data. In addition, we also infer lineages for B cells, Granulocyte cells, and Dendritic cells.

The dataset contains integrated embeddings, clustering results, and cell type annotations provided by the authors. Here we use CytoAnalyst to infer developmental trajectories given the provided embeddings and clusters. Specifically, we use the expression of four genes corresponding to four cell type lineages to infer the trajectories: Cd34 (HSPC), Ms4a1 (B cells), Ltf (Granulocyte), Siglech (Dendritic cells). Supplementary Figures S8, S11, S12, and S13 show the expression of these genes in the dataset, side-by-side with the clustering results. For each gene, we examine the expression pattern across clusters and manually select the start and end points for the trajectory inference. Supplementary Figure S9 shows the parameter settings we use for trajectory inference including embeddings, start groups, end groups, distance method, convergence threshold, etc. Supplementary Figures S10, S14, S15, and S16 show the inferred trajectories using CytoAnalyst. We also compare our results against the results reported by the authors. Supplementary Figure S17 shows that the results from CytoAnalyst match the published results, demonstrating the platform’s ability to reproduce complex trajectory inference analyses. Supplementary Section 2 provides details for the complete analysis workflow and results.

## 5 Conclusions

In this article, we present CytoAnalyst, a web-based platform for single-cell data analysis that is both powerful and easily accessible. By combining state-of-the-art analytical methods with advanced visualization techniques and an intuitive user interface, the platform enables researchers of all backgrounds to perform rigorous and comprehensive single-cell analysis. CytosAnalyst’s comprehensive analysis workflow includes data filtering, quality control, multi-sample integration, dimensionality reduction, cluster analysis, markers identification through DE analysis, cell type annotation, and pseudo-time trajectory inference. These analytical modules are interconnected with interactive visualization and systematic study management, enabling seamless transitions between steps while maintaining full parameter customization capabilities.

Several aspects distinguish CytoAnalyst from existing tools for single-cell analysis. First, its advanced visualization framework with multiple blending modes and interactive selection capabilities facilitates deep exploration of cellular heterogeneity. Second, the platform’s robust annotation system, powered by extensive reference databases and machine learning approaches, enables reliable cell type identification. Third, the implementation of real-time collaboration features and comprehensive project sharing capabilities promotes team-based analysis and reproducible research. CytoAnalyst’s scalable architecture and high-performance computing capabilities ensure it can handle the growing scale and complexity of single-cell datasets.

As single-cell technologies continue advancing, CytoAnalyst will evolve to incorporate new analytical methods while maintaining its core mission of democratizing single-cell transcriptomics analysis.

## 6 Data availability

CytoAnalyst is freely available at https://cytoanalyst.tinnguyen-lab.com and requires no registration or installation. The datasets used in the case studies are available at:

- https://cellxgene.cziscience.com/collections/59cd85c5-3b22-4035-b628-2a20810ad54b
- https://cellxgene.cziscience.com/collections/c353707f-09a4-4f12-92a0-cb741e57e5f0
- https://export.uppmax.uu.se/naiss2023-23-3/workshops/workshop-scrnaseq/trajectory/trajectory_seurat_filtered.h5ad

## 7 Additional files

**SupplementaryNote.pdf** : A text document describing the details of case studies 2 and 3.

**Supplementary-Table-S1.xlsx**: A spreadsheet containing differential expression analysis results for identifying general cluster markers of case study 2.

**Supplementary-Table-S2.xlsx**: A spreadsheet containing differential expression analysis results for identifying fibroblast-specific markers in young samples of case study 2.

**Supplementary-Table-S3.xlsx**: A spreadsheet containing differential expression analysis results for identifying fibroblast-specific markers in old samples of case study 2.

**Supplementary-Table-S4.xlsx**: A spreadsheet containing differential expression analysis results for comparing fibroblast subtypes in young samples of case study 2.

## 8 Funding

This work was partially supported by NSF (2343019 and 2203236), NIGMS (R44GM152152), and NCI (U01CA274573). Any opinions, findings, and conclusions or recommendations expressed in this material are those of the authors and do not necessarily reflect the views of any of the funding agencies.

## Notes

### Competing Interest Statement

The authors have declared no competing interest.

## References

[1] Malte D Luecken and Fabian J Theis. Current best practices in single-cell rna-seq analysis: a tutorial. Molecular Systems Biology, 15(6):e8746, 2019.

[2] Luke Zappia, Belinda Phipson, and Alicia Oshlack. Exploring the single-cell RNA-seq analysis landscape with the scRNA-tools database. PLoS Computational Biology, 14(6):e1006245, 2018.

[3] Andrew Butler, Paul Hoffman, Peter Smibert, Efthymia Papalexi, and Rahul Satija. Integrating single-cell transcriptomic data across different conditions, technologies, and species. Nature Biotechnology, 36:411–420, 2018.

[4] F. Alexander Wolf, Philipp Angerer, and Fabian J. Theis. SCANPY: large-scale single-cell gene expression data analysis. Genome Biology, 19:15, 2018.

[5] Cole Trapnell, Davide Cacchiarelli, Jonna Grimsby, Prapti Pokharel, Shuqiang Li, Michael Morse, Niall J Lennon, Kenneth J Livak, Tarjei S Mikkelsen, and John L Rinn. The dynamics and regulators of cell fate decisions are revealed by pseudotemporal ordering of single cells. Nature Biotechnology, 32:381–386, 2014.

[6] Adam Gayoso, Romain Lopez, Galen Xing, Pierre Boyeau, Valeh Valiollah Pour Amiri, Justin Hong, Katherine Wu, Michael Jayasuriya, Edouard Mehlman, Maxime Langevin, Yining Liu, Jules Samaran, Gabriel Misrachi, Achille Nazaret, Oscar Clivio, Chenling Xu, Tal Ashuach, Mariano Gabitto, Mohammad Lotfollahi, Valentine Svensson, Eduardo da Veiga Beltrame, Vitalii Kleshchevnikov, Carlos Talavera-López, Lior Pachter, Fabian J. Theis, Aaron Streets, Michael I. Jordan, Jeffrey Regier, and Nir Yosef. A Python library for probabilistic analysis of single-cell omics data. Nature Biotechnology, 40(2):163–166, 2022.

[7] Kevin Rue-Albrecht, Federico Marini, Charlotte Soneson, and Aaron TL Lun. iSEE: interactive summarizedexperiment explorer. F1000Research, 7:741, 2018.

[8] John F Ouyang, Uma S Kamaraj, Elaine Y Cao, and Owen JL Rackham. ShinyCell: simple and sharable visualization of single-cell gene expression data. Bioinformatics, 37(19):3374–3376, 2021.

[9] 10x Genomics. 10x Genomics Loupe Browser v8.0.0. https://www.10xgenomics.com/software, 2020.

[10] Biomage Ltd. Cellenics. https://www.biomage.net, 2022.

[11] Wendell J Pereira, Felipe Marques Almeida, D Conde, KM Balmant, PM Triozzi, HW Schmidt, C Dervinis, GJ Pappas, and M Kirst. Asc-Seurat: analytical single-cell Seurat-based web application. BMC Bioinformatics, 22:1–14, 2021.

[12] Ayman Yousif, Nizar Drou, Jillian Rowe, Mohammed Khalfan, and Kristin C Gunsalus. NASQAR: a web-based platform for high-throughput sequencing data analysis and visualization. BMC Bioinformatics, 21:1–14, 2020.

[13] Yichen Wang, Irzam Sarfraz, Nida Pervaiz, Rui Hong, Yusuke Koga, Vidya Akavoor, Xinyun Cao, Salam Alabdullatif, Syed Ali Zaib, Zhe Wang, Frederick Jansen, Masanao Yajima, W. Evan Johnson, and Joshua D. Campbell. Interactive analysis of single-cell data using flexible workflows with SCTK2. Patterns, 4(8), 2023.

[14] Shibla CZI Cell Science Program and Abdulla, Brian Aevermann, Pedro Assis, Seve Badajoz, Sidney M Bell, Emanuele Bezzi, Batuhan Cakir, Jim Chaffer, Signe Chambers, J Michael Cherry, Tiffany Chi, Jennifer Chien, Leah Dorman, Pablo Garcia-Nieto, Nayib Gloria, Mim Hastie, Daniel Hegeman, Jason Hilton, Timmy Huang, Amanda Infeld, Ana-Maria Istrate, Ivana Jelic, Kuni Katsuya, Yang Joon Kim, Karen Liang, Mike Lin, Maximilian Lombardo, Bailey Marshall, Bruce Martin, Fran McDade, Colin Megill, Nikhil Patel, Alexander Predeus, Brian Raymor, Behnam Robatmili, Dave Rogers, Erica Rutherford, Dana Sadgat, Andrew Shin, Corinn Small, Trent Smith, Prathap Sridharan, Alexander Tarashansky, Norbert Tavares, Harley Thomas, Andrew Tolopko, Meghan Urisko, Joyce Yan, Garabet Yeretssian, Jennifer Zamanian, Arathi Mani, Jonah Cool, and Ambrose Carr. CZ CELLxGENE discover: a single-cell data platform for scalable exploration, analysis and modeling of aggregated data. Nucleic Acids Research, 53(D1):D886–D900, 2025.

[15] Caleb Weinreb, Samuel Wolock, and Allon M Klein. SPRING: a kinetic interface for visualizing high dimensional single-cell expression data. Bioinformatics, 34(7):1246–1248, 2018.

[16] Kristofer Davie, Jasper Janssens, Duygu Koldere, Maxime De Waegeneer, Uli Pech, Lukasz Kreft, Sara Aibar, Samira Makhzami, Valerie Christiaens, Carmen Bravo González-Blas, Suresh Poovathingal, Gert Hulselmans, Katina I. Spanier, Thomas Moerman, Bram Vanspauwen, Sarah Geurs, Thierry Voet, Jeroen Lammertyn, Bernard Thienpont, Sha Liu, Nikos Konstantinides, Mark Fiers, Patrik Verstreken, and Stein Aerts. A Single-Cell Transcriptome Atlas of the Aging Drosophila Brain. Cell, 174(4):982–998, 2018.

[17] Vincent Gardeux, Fabrice PA David, Adrian Shajkofci, Petra C Schwalie, and Bart Deplancke. ASAP: a web-based platform for the analysis and interactive visualization of single-cell RNA-seq data. Bioinformatics, 33(19):3123–3125, 2017.

[18] Euxhen Hasanaj, Jingtao Wang, Arjun Sarathi, Jun Ding, and Ziv Bar-Joseph. Interactive single-cell data analysis using Cellar. Nature Communications, 13(1):1998, 2022.

[19] Raman Sethi, Kok Siong Ang, Mengwei Li, Yahui Long, Jingjing Ling, and Jinmiao Chen. ezSingleCell: an integrated one-stop single-cell and spatial omics analysis platform for bench scientists. Nature Communications, 15(1):5600, 2024.

[20] Matthew L Speir, Aparna Bhaduri, Nikolay S Markov, Pablo Moreno, Tomasz J Nowakowski, Irene Papatheodorou, Alex A Pollen, Brian J Raney, Lucas Seninge, W James Kent, and Maximilian Haeussler. UCSC Cell Browser: visualize your single-cell data. Bioinformatics, 37(23):4578–4580, 2021.

[21] Andrew Jiang, Klaus Lehnert, Linya You, and Russell G Snell. ICARUS, an interactive web server for single cell RNA-seq analysis. Nucleic Acids Research, 50(W1):W427–W433, 2022.

[22] Carlos Prieto, David Barrios, and Angela Villaverde. SingleCAnalyzer: interactive analysis of single cell RNA-Seq data on the cloud. Frontiers in Bioinformatics, 2:793309, 2022.

[23] Amir Alavi, Matthew Ruffalo, Aiyappa Parvangada, Zhilin Huang, and Ziv Bar-Joseph. A web server for comparative analysis of single-cell RNA-seq data. Nature Communications, 9(1):4768, 2018.

[24] Fei Quan, Xin Liang, Mingjiang Cheng, Huan Yang, Kun Liu, Shengyuan He, Shangqin Sun, Menglan Deng, Yanzhen He, Wei Liu, Shuai Wang, Shuxiang Zhao, Lantian Deng, Xiaobo Hou, Xinxin Zhang, and Yun Xiao. Annotation of cell types (ACT): a convenient web server for cell type annotation. Genome Medicine, 15(1):91, 2023.

[25] Weideng Wei, Xiaoqiang Xia, Taiwen Li, Qianming Chen, and Xiaodong Feng. Shaoxia: a web-based interactive analysis platform for single cell RNA sequencing data. BMC Genomics, 25(1):402, 2024.

[26] Andrew P Voigt, S Scott Whitmore, Nicholas D Lessing, Adam P DeLuca, Budd A Tucker, Edwin M Stone, Robert F Mullins, and Todd E Scheetz. Spectacle: an interactive resource for ocular single-cell RNA sequencing data analysis. Experimental Eye Research, 200:108204, 2020.

[27] Hao Yu, Yuqing Wang, Xi Zhang, and Zheng Wang. GRACE: a comprehensive web-based platform for integrative single-cell transcriptome analysis. NAR Genomics and Bioinformatics, 5(2):lqad050, 2023.

[28] Oscar Franzén and Johan LM Björkegren. alona: a web server for single-cell RNA-seq analysis. Bioinformatics, 36(12):3910–3912, 2020.

[29] Xun Zhu, Thomas K Wolfgruber, Austin Tasato, Cédric Arisdakessian, David G Garmire, and Lana X Garmire. Granatum: a graphical single-cell RNA-Seq analysis pipeline for genomics scientists. Genome Medicine, 9:1–12, 2017.

[30] Grace X. Y. Zheng, Jessica M. Terry, Phillip Belgrader, Paul Ryvkin, Zachary W. Bent, Ryan Wilson, Solongo B. Ziraldo, Tobias D. Wheeler, Geoff P. McDermott, Junjie Zhu, Mark T. Gregory, Joe Shuga, Luz Montesclaros, Jason G. Underwood, Donald A. Masquelier, Stefanie Y. Nishimura, Michael Schnall-Levin, Paul W. Wyatt, Christopher M. Hindson, Rajiv Bharadwaj, Alexander Wong, Kevin D. Ness, Lan W. Beppu, H. Joachim Deeg, Christopher McFarland, Keith R. Loeb, William J. Valente, Nolan G. Ericson, Emily A. Stevens, Jerald P. Radich, Tarjei S. Mikkelsen, Benjamin J. Hindson, and Jason H. Bielas. Massively parallel digital transcriptional profiling of single cells. Nature Communications, 8:14049, Jan 2017.

[31] Isaac Virshup, Sergei Rybakov, Fabian J Theis, Philipp Angerer, and F Alexander Wolf. anndata: Access and store annotated data matrices. Journal of Open Source Software, 9(101):4371, 2024.

[32] Laurens Van der Maaten and Geoffrey Hinton. Visualizing data using t-SNE. Journal of Machine Learning Research, 9(11), 2008.

[33] Leland McInnes, John Healy, and James Melville. Umap: uniform manifold approximation and projection for dimension reduction. *arXiv preprint arXiv:1802.03426*, 2018.

[34] Tim Stuart, Andrew Butler, Paul Hoffman, Christoph Hafemeister, Efthymia Papalexi, William M Mauck III, Yuhan Hao, Marlon Stoeckius, Peter Smibert, and Rahul Satija. Comprehensive integration of single-cell data. Cell, 177(7):1888–1902, 2019.

[35] Christoph Hafemeister and Rahul Satija. Normalization and variance stabilization of single-cell RNA-seq data using regularized negative binomial regression. Genome Biology, 20(1):296, 2019.

[36] Blythe Durbin, Johanna Hardin, Douglas Hawkins, and David Rocke. A variance-stabilizing transformation for gene-expression microarray data. Bioinformatics, 18, 2002.

[37] Joseph B Patlak. Measuring kinetics of complex single ion channel data using mean-variance histograms. Biophysical Journal, 65(1):29–42, 1993.

[38] Yuhan Hao, Stephanie Hao, Erica Andersen-Nissen, William M. Mauck, Shiwei Zheng, Andrew Butler, Maddie J. Lee, Aaron J. Wilk, Charlotte Darby, Michael Zager, Paul Hoffman, Marlon Stoeckius, Efthymia Papalexi, Eleni P. Mimitou, Jaison Jain, Avi Srivastava, Tim Stuart, Lamar M. Fleming, Bertrand Yeung, Angela J. Rogers, Juliana M. McElrath, Catherine A. Blish, Raphael Gottardo, Peter Smibert, and Rahul Satija. Integrated analysis of multimodal single-cell data. Cell, 184(13):3573–3587.e29, 2021.

[39] Yuhan Hao, Tim Stuart, Madeline H Kowalski, Saket Choudhary, Paul Hoffman, Austin Hartman, Avi Srivastava, Gesmira Molla, Shaista Madad, Carlos Fernandez-Granda, and Rahul Satija. Dictionary learning for integrative, multimodal and scalable single-cell analysis. Nature Biotechnology, 42(2):293– 304, 2024.

[40] Ilya Korsunsky, Nghia Millard, Jean Fan, Kamil Slowikowski, Fan Zhang, Kevin Wei, Yuriy Baglaenko, Michael Brenner, Po-ru Loh, and Soumya Raychaudhuri. Fast, sensitive and accurate integration of single-cell data with harmony. Nature Methods, 16(12):1289–1296, 2019.

[41] Hervé Abdi and Lynne J Williams. Principal component analysis. Wiley Interdisciplinary Reviews: Computational Statistics, 2(4):433–459, 2010.

[42] Vincent D Blondel, Jean-Loup Guillaume, Renaud Lambiotte, and Etienne Lefebvre. Fast unfolding of communities in large networks. Journal of Statistical Mechanics: Theory and Experiment, 2008(10):P10008, 2008.

[43] Vincent A. Traag, Ludo Waltman, and Nees Jan van Eck. From Louvain to Leiden: guaranteeing well-connected communities. Scientific Reports, 9(1):1–12, 2019.

[44] Trupti Kodinariya and Prashant Makwana. Review on determining number of Cluster in K-Means Clustering. International Journal of Advance Research in Computer Science and Management Studies, 1:90–95, 01 2013.

[45] Shixiong Zhang, Xiangtao Li, Jiecong Lin, Qiuzhen Lin, and Ka-Chun Wong. Review of single-cell RNA-seq data clustering for cell-type identification and characterization. RNA, 29(5):517–530, 2023.

[46] Frank Wilcoxon, SK Katti, and Roberta A Wilcox. Critical values and probability levels for the Wilcoxon rank sum test and the Wilcoxon signed rank test. Selected tables in mathematical statistics, 1:171–259, 1970.

[47] Greg Finak, Andrew McDavid, Masanao Yajima, Jingyuan Deng, Vivian Gersuk, Alex K. Shalek, Chloe K. Slichter, Hannah W. Miller, M. Juliana McElrath, Martin Prlic, Peter S. Linsley, and Raphael Gottardo. MAST: a flexible statistical framework for assessing transcriptional changes and characterizing heterogeneity in single-cell RNA sequencing data. Genome Biology, 16:278, 2015.

[48] David Thissen, Lynne Steinberg, and Daniel Kuang. Quick and easy implementation of the BenjaminiHochberg procedure for controlling the false positive rate in multiple comparisons. Journal of Educational and Behavioral Statistics, 27(1):77–83, 2002.

[49] Congxue Hu, Tengyue Li, Yingqi Xu, Xinxin Zhang, Feng Li, Jing Bai, Jing Chen, Wenqi Jiang, Kaiyue Yang, Qi Ou, Xia Li, Peng Wang, and Yunpeng Zhang. CellMarker 2.0: an updated database of manually curated cell markers in human/mouse and web tools based on scRNA-seq data. Nucleic Acids Research, 51(D1):D870–D876, 2023.

[50] Aaron Grattafiori, Abhimanyu Dubey, Abhinav Jauhri, Abhinav Pandey, Abhishek Kadian, Ahmad Al-Dahle, Aiesha Letman, Akhil Mathur, Alan Schelten, Alex Vaughan, Amy Yang, Angela Fan, Anirudh Goyal, Anthony Hartshorn, Aobo Yang, Archi Mitra, Archie Sravankumar, Artem Korenev, Arthur Hinsvark, Arun Rao, Aston Zhang, Aurelien Rodriguez, Austen Gregerson, Ava Spataru, Baptiste Roziere, Bethany Biron, Binh Tang, Bobbie Chern, Charlotte Caucheteux, Chaya Nayak, Chloe Bi, Chris Marra, Chris McConnell, Christian Keller, Christophe Touret, Chunyang Wu, Corinne Wong, Cristian Canton Ferrer, Cyrus Nikolaidis, Damien Allonsius, Daniel Song, Danielle Pintz, Danny Livshits, Danny Wyatt, David Esiobu, Dhruv Choudhary, Dhruv Mahajan, Diego Garcia-Olano, Diego Perino, Dieuwke Hupkes, Egor Lakomkin, Ehab AlBadawy, Elina Lobanova, Emily Dinan, Eric Michael Smith, Filip Radenovic, Francisco Guzmán, Frank Zhang, Gabriel Synnaeve, Gabrielle Lee, Georgia Lewis Anderson, Govind Thattai, Graeme Nail, Gregoire Mialon, Guan Pang, Guillem Cucurell, Hailey Nguyen, Hannah Korevaar, Hu Xu, Hugo Touvron, Iliyan Zarov, Imanol Arrieta Ibarra, Isabel Kloumann, Ishan Misra, Ivan Evtimov, Jack Zhang, Jade Copet, Jaewon Lee, Jan Geffert, Jana Vranes, Jason Park, Jay Mahadeokar, Jeet Shah, Jelmer van der Linde, Jennifer Billock, Jenny Hong, Jenya Lee, Jeremy Fu, Jianfeng Chi, Jianyu Huang, Jiawen Liu, Jie Wang, Jiecao Yu, Joanna Bitton, Joe Spisak, Jongsoo Park, Joseph Rocca, Joshua Johnstun, Joshua Saxe, Junteng Jia, Kalyan Vasuden Alwala, Karthik Prasad, Kartikeya Upasani, Kate Plawiak, Ke Li, Kenneth Heafield, Kevin Stone, Khalid El-Arini, Krithika Iyer, Kshitiz Malik, Kuenley Chiu, Kunal Bhalla, Kushal Lakhotia, Lauren Rantala-Yeary, Laurens van der Maaten, Lawrence Chen, Liang Tan, Liz Jenkins, Louis Martin, Lovish Madaan, Lubo Malo, Lukas Blecher, Lukas Landzaat, Luke de Oliveira, Madeline Muzzi, Mahesh Pasupuleti, Mannat Singh, Manohar Paluri, Marcin Kardas, Maria Tsimpoukelli, Mathew Oldham, Mathieu Rita, Maya Pavlova, Melanie Kambadur, Mike Lewis, Min Si, Mitesh Kumar Singh, Mona Hassan, Naman Goyal, Narjes Torabi, Nikolay Bashlykov, Nikolay Bogoychev, Niladri Chatterji, Ning Zhang, Olivier Duchenne, Onur Çelebi, Patrick Alrassy, Pengchuan Zhang, Pengwei Li, Petar Vasic, Peter Weng, Prajjwal Bhargava, Pratik Dubal, Praveen Krishnan, Punit Singh Koura, Puxin Xu, Qing He, Qingxiao Dong, Ragavan Srinivasan, Raj Ganapathy, Ramon Calderer, Ricardo Silveira Cabral, Robert Stojnic, Roberta Raileanu, Rohan Maheswari, Rohit Girdhar, Rohit Patel, Romain Sauvestre, Ronnie Polidoro, Roshan Sumbaly, Ross Taylor, Ruan Silva, Rui Hou, Rui Wang, Saghar Hosseini, Sahana Chennabasappa, Sanjay Singh, Sean Bell, Seohyun Sonia Kim, Sergey Edunov, Shaoliang Nie, Sharan Narang, Sharath Raparthy, Sheng Shen, Shengye Wan, Shruti Bhosale, Shun Zhang, Simon Vandenhende, Soumya Batra, Spencer Whitman, Sten Sootla, Stephane Collot, Suchin Gururangan, Sydney Borodinsky, Tamar Herman, Tara Fowler, Tarek Sheasha, Thomas Georgiou, Thomas Scialom, Tobias Speckbacher, Todor Mihaylov, Tong Xiao, Ujjwal Karn, Vedanuj Goswami, Vibhor Gupta, Vignesh Ramanathan, Viktor Kerkez, Vincent Gonguet, Virginie Do, Vish Vogeti, Vítor Albiero, Vladan Petrovic, Weiwei Chu, Wenhan Xiong, Wenyin Fu, Whitney Meers, Xavier Martinet, Xiaodong Wang, Xiaofang Wang, Xiaoqing Ellen Tan, Xide Xia, Xinfeng Xie, Xuchao Jia, Xuewei Wang, Yaelle Goldschlag, Yashesh Gaur, Yasmine Babaei, Yi Wen, Yiwen Song, Yuchen Zhang, Yue Li, Yuning Mao, Zacharie Delpierre Coudert, Zheng Yan, Zhengxing Chen, Zoe Papakipos, Aaditya Singh, Aayushi Srivastava, Abha Jain, Adam Kelsey, Adam Shajnfeld, Adithya Gangidi, Adolfo Victoria, Ahuva Goldstand, Ajay Menon, Ajay Sharma, Alex Boesenberg, Alexei Baevski, Allie Feinstein, Amanda Kallet, Amit Sangani, Amos Teo, Anam Yunus, Andrei Lupu, Andres Alvarado, Andrew Caples, Andrew Gu, Andrew Ho, Andrew Poulton, Andrew Ryan, Ankit Ramchandani, Annie Dong, Annie Franco, Anuj Goyal, Aparajita Saraf, Arkabandhu Chowdhury, Ashley Gabriel, Ashwin Bharambe, Assaf Eisenman, Azadeh Yazdan, Beau James, Ben Maurer, Benjamin Leonhardi, Bernie Huang, Beth Loyd, Beto De Paola, Bhargavi Paranjape, Bing Liu, Bo Wu, Boyu Ni, Braden Hancock, Bram Wasti, Brandon Spence, Brani Stojkovic, Brian Gamido, Britt Montalvo, Carl Parker, Carly Burton, Catalina Mejia, Ce Liu, Changhan Wang, Changkyu Kim, Chao Zhou, Chester Hu, Ching-Hsiang Chu, Chris Cai, Chris Tindal, Christoph Feichtenhofer, Cynthia Gao, Damon Civin, Dana Beaty, Daniel Kreymer, Daniel Li, David Adkins, David Xu, Davide Testuggine, Delia David, Devi Parikh, Diana Liskovich, Didem Foss, Dingkang Wang, Duc Le, Dustin Holland, Edward Dowling, Eissa Jamil, Elaine Montgomery, Eleonora Presani, Emily Hahn, Emily Wood, Eric-Tuan Le, Erik Brinkman, Esteban Arcaute, Evan Dunbar, Evan Smothers, Fei Sun, Felix Kreuk, Feng Tian, Filippos Kokkinos, Firat Ozgenel, Francesco Caggioni, Frank Kanayet, Frank Seide, Gabriela Medina Florez, Gabriella Schwarz, Gada Badeer, Georgia Swee, Gil Halpern, Grant Herman, Grigory Sizov, Guangyi, Zhang, Guna Lakshminarayanan, Hakan Inan, Hamid Shojanazeri, Han Zou, Hannah Wang, Hanwen Zha, Haroun Habeeb, Harrison Rudolph, Helen Suk, Henry Aspegren, Hunter Goldman, Hongyuan Zhan, Ibrahim Damlaj, Igor Molybog, Igor Tufanov, Ilias Leontiadis, Irina-Elena Veliche, Itai Gat, Jake Weissman, James Geboski, James Kohli, Janice Lam, Japhet Asher, Jean-Baptiste Gaya, Jeff Marcus, Jeff Tang, Jennifer Chan, Jenny Zhen, Jeremy Reizenstein, Jeremy Teboul, Jessica Zhong, Jian Jin, Jingyi Yang, Joe Cummings, Jon Carvill, Jon Shepard, Jonathan McPhie, Jonathan Torres, Josh Ginsburg, Junjie Wang, Kai Wu, Kam Hou U, Karan Saxena, Kartikay Khandelwal, Katayoun Zand, Kathy Matosich, Kaushik Veeraraghavan, Kelly Michelena, Keqian Li, Kiran Jagadeesh, Kun Huang, Kunal Chawla, Kyle Huang, Lailin Chen, Lakshya Garg, Lavender A, Leandro Silva, Lee Bell, Lei Zhang, Liangpeng Guo, Licheng Yu, Liron Moshkovich, Luca Wehrstedt, Madian Khabsa, Manav Avalani, Manish Bhatt, Martynas Mankus, Matan Hasson, Matthew Lennie, Matthias Reso, Maxim Groshev, Maxim Naumov, Maya Lathi, Meghan Keneally, Miao Liu, Michael L. Seltzer, Michal Valko, Michelle Restrepo, Mihir Patel, Mik Vyatskov, Mikayel Samvelyan, Mike Clark, Mike Macey, Mike Wang, Miquel Jubert Hermoso, Mo Metanat, Mohammad Rastegari, Munish Bansal, Nandhini Santhanam, Natascha Parks, Natasha White, Navyata Bawa, Nayan Singhal, Nick Egebo, Nicolas Usunier, Nikhil Mehta, Nikolay Pavlovich Laptev, Ning Dong, Norman Cheng, Oleg Chernoguz, Olivia Hart, Omkar Salpekar, Ozlem Kalinli, Parkin Kent, Parth Parekh, Paul Saab, Pavan Balaji, Pedro Rittner, Philip Bontrager, Pierre Roux, Piotr Dollar, Polina Zvyagina, Prashant Ratanchandani, Pritish Yuvraj, Qian Liang, Rachad Alao, Rachel Rodriguez, Rafi Ayub, Raghotham Murthy, Raghu Nayani, Rahul Mitra, Rangaprabhu Parthasarathy, Raymond Li, Rebekkah Hogan, Robin Battey, Rocky Wang, Russ Howes, Ruty Rinott, Sachin Mehta, Sachin Siby, Sai Jayesh Bondu, Samyak Datta, Sara Chugh, Sara Hunt, Sargun Dhillon, Sasha Sidorov, Satadru Pan, Saurabh Mahajan, Saurabh Verma, Seiji Yamamoto, Sharadh Ramaswamy, Shaun Lindsay, Shaun Lindsay, Sheng Feng, Shenghao Lin, Shengxin Cindy Zha, Shishir Patil, Shiva Shankar, Shuqiang Zhang, Shuqiang Zhang, Sinong Wang, Sneha Agarwal, Soji Sajuyigbe, Soumith Chintala, Stephanie Max, Stephen Chen, Steve Kehoe, Steve Satterfield, Sudarshan Govindaprasad, Sumit Gupta, Summer Deng, Sungmin Cho, Sunny Virk, Suraj Subramanian, Sy Choudhury, Sydney Goldman, Tal Remez, Tamar Glaser, Tamara Best, Thilo Koehler, Thomas Robinson, Tianhe Li, Tianjun Zhang, Tim Matthews, Timothy Chou, Tzook Shaked, Varun Vontimitta, Victoria Ajayi, Victoria Montanez, Vijai Mohan, Vinay Satish Kumar, Vishal Mangla, Vlad Ionescu, Vlad Poenaru, Vlad Tiberiu Mihailescu, Vladimir Ivanov, Wei Li, Wenchen Wang, Wenwen Jiang, Wes Bouaziz, Will Constable, Xiaocheng Tang, Xiaojian Wu, Xiaolan Wang, Xilun Wu, Xinbo Gao, Yaniv Kleinman, Yanjun Chen, Ye Hu, Ye Jia, Ye Qi, Yenda Li, Yilin Zhang, Ying Zhang, Yossi Adi, Youngjin Nam, Yu, Wang, Yu Zhao, Yuchen Hao, Yundi Qian, Yunlu Li, Yuzi He, Zach Rait, Zachary DeVito, Zef Rosnbrick, Zhaoduo Wen, Zhenyu Yang, Zhiwei Zhao, and Zhiyu Ma. The llama 3 herd of models. arXiv preprint arXiv:2407.21783, 2024.

[51] Bernard L. Welch. The generalization of “Student’s” problem when several different population varlances are involved. Biometrika, 34(1-2):28–35, 1947.

[52] Gennady Korotkevich, Vladimir Sukhov, Nikolay Budin, Boris Shpak, Maxim N Artyomov, and Alexey Sergushichev. Fast gene set enrichment analysis. bioRxiv, page 060012, 2016.

[53] Kelly Street, Davide Risso, Russell B Fletcher, Diya Das, John Ngai, Nir Yosef, Elizabeth Purdom, and Sandrine Dudoit. Slingshot: cell lineage and pseudotime inference for single-cell transcriptomics. BMC Genomics, 19:477, 2018.

[54] Wouter Saelens, Robrecht Cannoodt, Helena Todorov, and Yvan Saeys. A comparison of single-cell trajectory inference methods. Nature Biotechnology, 37:547–554, 2019.

[55] Trevor Hastie and Werner Stuetzle. Principal curves. Journal of the American Statistical Association, 84(406):502–516, 1989.

[56] Stephanie Frenz-Wiessner, Savannah D. Fairley, Maximilian Buser, Isabel Goek, Kirill Salewskij, Gustav Jonsson, David Illig, Benedicta zu Putlitz, Daniel Petersheim, Yue Li, Pin-Hsuan Chen, Martina Kalauz, Raffaele Conca, Michael Sterr, Johanna Geuder, Yoko Mizoguchi, Remco T. A. Megens, Monika I. Linder, Daniel Kotlarz, Martina Rudelius, Josef M. Penninger, Carsten Marr, and Christoph Klein. Generation of complex bone marrow organoids from human induced pluripotent stem cells. Nature Methods, 21:868–881, 2024.

[57] Llorenç Solé-Boldo, Günter Raddatz, Sabrina Schütz, Jan-Philipp Mallm, Karsten Rippe, Anke S Lonsdorf, Manuel Rodríguez-Paredes, and Frank Lyko. Single-cell transcriptomes of the human skin reveal age-related loss of fibroblast priming. Communications Biology, 3(1):188, 2020.

[58] Xiaoping Han, Renying Wang, Yincong Zhou, Lijiang Fei, Huiyu Sun, Shujing Lai, Assieh Saadatpour, Ziming Zhou, Haide Chen, Fang Ye, Daosheng Huang, Yang Xu, Wentao Huang, Mengmeng Jiang, Xinyi Jiang, Jie Mao, Yao Chen, Chenyu Lu, Jin Xie, Qun Fang, Yibin Wang, Rui Yue, Tiefeng Li, He Huang, Stuart H. Orkin, Guo-Cheng Yuan, Ming Chen, and Guoji Guo. Mapping the Mouse Cell Atlas by Microwell-Se. Cell, 172(5):1091–1107, 2018.

[59] Nicholas Schaum, Jim Karkanias, Norma F. Neff, Andrew P. May, Stephen R. Quake, Tony Wyss-Coray, Spyros Darmanis, Joshua Batson, Olga Botvinnik, Michelle B. Chen, Steven Chen, Foad Green, Robert C. Jones, Ashley Maynard, Lolita Penland, Angela Oliveira Pisco, Rene V. Sit, Geoffrey M. Stanley, James T. Webber, Fabio Zanini, Ankit S. Baghel, Isaac Bakerman, Ishita Bansal, Daniela Berdnik, Biter Bilen, Douglas Brownfield, Corey Cain, Michelle B. Chen, Steven Chen, Min Cho, Giana Cirolia, Stephanie D. Conley, Spyros Darmanis, Aaron Demers, Kubilay Demir, Antoine de Morree, Tessa Divita, Haley du Bois, Laughing Bear Torrez Dulgeroff, Hamid Ebadi, F. Hernán Espinoza, Matt Fish, Qiang Gan, Benson M. George, Astrid Gillich, Foad Green, Geraldine Genetiano, Xueying Gu, Gunsagar S. Gulati, Yan Hang, Shayan Hosseinzadeh, Albin Huang, Tal Iram, Taichi Isobe, Feather Ives, Robert C. Jones, Kevin S. Kao, Guruswamy Karnam, Aaron M. Kershner, Bernhard M. Kiss, William Kong, Maya E. Kumar, Jonathan Y. Lam, Davis P. Lee, Song E. Lee, Guang Li, Qingyun Li, Ling Liu, Annie Lo, Wan-Jin Lu, Anoop Manjunath, Andrew P. May, Kaia L. May, Oliver L. May, Ashley Maynard, Marina McKay, Ross J. Metzger, Marco Mignardi, Dullei Min, Ahmad N. Nabhan, Norma F. Neff, Katharine M. Ng, Joseph Noh, Rasika Patkar, Weng Chuan Peng, Lolita Penland, Robert Puccinelli, Eric J. Rulifson, Nicholas Schaum, Shaheen S. Sikandar, Rahul Sinha, Rene V. Sit, Krzysztof Szade, Weilun Tan, Cristina Tato, Krissie Tellez, Kyle J. Travaglini, Carolina Tropini, Lucas Waldburger, Linda J. van Weele, Michael N. Wosczyna, Jinyi Xiang, Soso Xue, Justin Youngyunpipatkul, Fabio Zanini, Macy E. Zardeneta, Fan Zhang, Lu Zhou, Ishita Bansal, Steven Chen, Min Cho, Giana Cirolia, Spyros Darmanis, Aaron Demers, Tessa Divita, Hamid Ebadi, Geraldine Genetiano, Foad Green, Shayan Hosseinzadeh, Feather Ives, Annie Lo, Andrew P. May, Ashley Maynard, Marina McKay, Norma F. Neff, Lolita Penland, Rene V. Sit, Weilun Tan, Lucas Waldburger, Justin Youngyunpipatkul, Joshua Batson, Olga Botvinnik, Paola Castro, Derek Croote, Spyros Darmanis, Joseph L. DeRisi, Jim Karkanias, Angela Oliveira Pisco, Geoffrey M. Stanley, James T. Webber, Fabio Zanini, Ankit S. Baghel, Isaac Bakerman, Joshua Batson, Biter Bilen, Olga Botvinnik, Douglas Brownfield, Michelle B. Chen, Spyros Darmanis, Kubilay Demir, Antoine de Morree, Hamid Ebadi, F. Hernán Espinoza, Matt Fish, Qiang Gan, Benson M. George, Astrid Gillich, Xueying Gu, Gunsagar S. Gulati, Yan Hang, Albin Huang, Tal Iram, Taichi Isobe, Guruswamy Karnam, Aaron M. Kershner, Bernhard M. Kiss, William Kong, Christin S. Kuo, Jonathan Y. Lam, Benoit Lehallier, Guang Li, Qingyun Li, Ling Liu, Wan-Jin Lu, Dullei Min, Ahmad N. Nabhan, Katharine M. Ng, Patricia K. Nguyen, Rasika Patkar, Weng Chuan Peng, Lolita Penland, Eric J. Rulifson, Nicholas Schaum, Shaheen S. Sikandar, Rahul Sinha, Krzysztof Szade, Serena Y. Tan, Krissie Tellez, Kyle J. Travaglini, Carolina Tropini, Linda J. van Weele, Bruce M. Wang, Michael N. Wosczyna, Jinyi Xiang, Hanadie Yousef, Lu Zhou, Joshua Batson, Olga Botvinnik, Steven Chen, Spyros Darmanis, Foad Green, Andrew P. May, Ashley Maynard, Angela Oliveira Pisco, Stephen R. Quake, Nicholas Schaum, Geoffrey M. Stanley, James T. Webber, Tony Wyss-Coray, Fabio Zanini, Philip A. Beachy, Charles K. F. Chan, Antoine de Morree, Benson M. George, Gunsagar S. Gulati, Yan Hang, Kerwyn Casey Huang, Tal Iram, Taichi Isobe, Aaron M. Kershner, Bernhard M. Kiss, William Kong, Guang Li, Qingyun Li, Ling Liu, Wan-Jin Lu, Ahmad N. Nabhan, Katharine M. Ng, Patricia K. Nguyen, Weng Chuan Peng, Eric J. Rulifson, Nicholas Schaum, Shaheen S. Sikandar, Rahul Sinha, Krzysztof Szade, Kyle J. Travaglini, Carolina Tropini, Bruce M. Wang, Kenneth Weinberg, Michael N. Wosczyna, Sean M. Wu, Hanadie Yousef, Ben A. Barres, Philip A. Beachy, Charles K. F. Chan, Michael F. Clarke, Spyros Darmanis, Kerwyn Casey Huang, Jim Karkanias, Seung K. Kim, Mark A. Krasnow, Maya E. Kumar, Christin S. Kuo, Andrew P. May, Ross J. Metzger, Norma F. Neff, Roel Nusse, Patricia K. Nguyen, Thomas A. Rando, Justin Sonnenburg, Bruce M. Wang, Kenneth Weinberg, Irving L. Weissman, Sean M. Wu, Stephen R. Quake, Tony Wyss-Coray, The Tabula Muris Consortium, Overall coordination, Logistical coordination, Organ collection and processing, Library preparation and sequencing, Computational data analysis, Cell type annotation, Writing group, Supplemental text writing group, and Principal investigators. Single-cell transcriptomics of 20 mouse organs creates a Tabula Muris. Nature, 562(7727):367–372, 2018.

[60] Joakim S. Dahlin, Fiona K. Hamey, Blanca Pijuan-Sala, Mairi Shepherd, Winnie WY Lau, Sonia Nestorowa, Caleb Weinreb, Samuel Wolock, Rebecca Hannah, Evangelia Diamanti, David G. Kent, Berthold Göttgens, and Nicola K. Wilson. A single-cell hematopoietic landscape resolves 8 lineage trajectories and defects in Kit mutant mice. Blood, The Journal of the American Society of Hematology, 131(21):e1–e11, 2018.

[61] Samuel L Wolock, Romain Lopez, and Allon M Klein. Scrublet: computational identification of cell doublets in single-cell transcriptomic data. Cell Systems, 8(4):281–291, 2019.

[62] Åsa Björklund, Paulo Czarnewski, Susanne Reinsbach, and Roy Francis. Trajectory inference using Slingshot. https://nbisweden.github.io/workshop-scRNAseq/labs/seurat/seurat_07_trajectory.html, 2025. NBIS Workshop on Single-cell RNA-seq Analysis.

[63] Koen Van den Berge, Hector Roux de Bézieux, Kelly Street, Wouter Saelens, Robrecht Cannoodt, Yvan Saeys, Sandrine Dudoit, and Lieven Clement. Trajectory-based differential expression analysis for single-cell sequencing data. Nature Communications, 11(1):1201, 2020.

