## Supplementary Note for "CytoAnalyst: A web-based platform for comprehensive single-cell RNA sequencing analysis"

#### Supplementary Material

**Phi Bya [1], Duy Tran [1], Khoi Nguyen [1], Sorin Draghici [2], Tin Nguyen [1]**
[1]Department of Computer Science and Software Engineering, Auburn University, Auburn, AL 36849, USA
[2]Department of Computer Science, Wayne State University, Detroit, MI 48202, USA

### 1 Case Study 2: Cell Type Markers of Skin Samples

In this case study, we perform differential expression analyses on a dataset from Solé-Boldo et al. [1] using the CytoAnalyst platform. The authors collected cells from the sun-protected inguinoiliac region of whole-skin samples from five male donors: two young donors (25 and 27 years old) and three old donors (53, 69, and 70 years old). They processed the raw sequencing data using Cell Ranger v2.1.0 and conducted further analysis using Seurat package v3.1.1 in R v3.5.1. To ensure high-quality data, they removed potential doublets (cells with >7,500 expressed genes) and apoptotic cells (cells with >5% mitochondrial reads), resulting in 15,457 high-quality single-cell transcriptomes.

The authors integrated the dataset using Seurat's standard integration protocol by log-normalizing UMI counts, identifying 2,000 variable genes per sample, and using canonical correlation analysis (CCA) with 30 dimensions to integrate the data. They scaled the integrated data and performed principal component analysis (PCA) with 20 dimensions to construct a Shared Nearest Neighbor (SNN) Graph. Next, they used Louvain clustering with a resolution of 0.4 to identify 17 distinct clusters. They determined markers for each cluster by performing differential expression analysis using the Wilcoxon rank sum test, comparing gene expression of cells within each cluster against all other cells. By comparing known markers with the most representative expressed genes of each cluster, they identified 9 main cell types in the skin: keratinocytes, fibroblasts, macrophages/dendritic cells, T cells, vascular and lymphatic endothelial cells, pericytes, erythrocytes, and melanocytes.

For a deeper analysis of fibroblast heterogeneity, the authors isolated 5,948 fibroblasts (clusters 1, 2, 3, and 9) and performed functional enrichment to identify fibroblast subpopulations. The analysis first identified markers for each fibroblast subpopulation by performing differential expression analysis using the Wilcoxon rank sum test using only cells from young samples. They then performed gene ontology (GO) analysis using the DAVID Bioinformatics Database. Based on the enriched GO terms associated with each subpopulation, they identified and functionally characterized four main fibroblast subpopulations: secretory-reticular fibroblasts, pro-inflammatory fibroblasts, secretory-papillary fibroblasts, and mesenchymal fibroblasts. To confirm the findings, using cells from the four fibroblast subpopulations from young samples, they performed a second-level clustering analysis using the Louvain algorithm with 20 PCA dimensions and a resolution of 0.5. This analysis further divided the pro-inflammatory fibroblasts into two distinct clusters, pro-inflammatory fibroblasts A and B, resulting in a total of five fibroblast subpopulations. Finally, they validated these subpopulations using RNA fluorescence in situ hybridization (RNA FISH) and immunohistochemistry on independent skin sections.

The authors then investigated age-related changes in fibroblast transcriptomes by comparing fibroblasts from young and old donors. Then they performed differential expression analysis using the Wilcoxon rank sum test to identify genes differentially expressed with age in each fibroblast subpopulation. Their analysis revealed significant changes in cell cycle phase distribution, with increased G1/S phase delay in aged cells. They observed a loss of functional priming in aged fibroblasts, demonstrated by fewer function-related GO terms and decreased collagen gene expression. The authors also detected changes in papillary and reticular gene signatures and identified subpopulation-specific expression of skin aging-associated secreted proteins (SAASP). Finally, using CellPhoneDB v2.0.0, they analyzed cell-cell interactions and discovered a substantial reduction in predicted interactions between aged dermal fibroblasts and other skin cells, particularly undifferentiated keratinocytes at the dermal-epidermal junction.

In this analysis, we aim to reproduce the differential expression analysis results from the original study and evaluate the performance of CytoAnalyst. Specifically, we conduct four distinct analyses: (1) identification of general cluster markers (all clusters), (2) Fibroblast-specific markers in young samples (clusters 1, 2, 3, and 9), (3) Fibroblast-specific markers in old samples (clusters 1, 2, 3, and 9), and (4) detailed subpopulation analysis of Fibroblasts in young samples using second-level clustering. Overall, CytoAnalyst's results align closely with the original study, with the majority of differentially expressed genes identified in the original study also detected by CytoAnalyst.

#### 1.1 DE Analysis to Indentify Markers of 17 Clusters

In this analysis, we utilize 17 clusters from the original study and compare each cluster against all other clusters to identify their markers. The analysis settings in CytoAnalyst's interface are shown in Figure S1A.

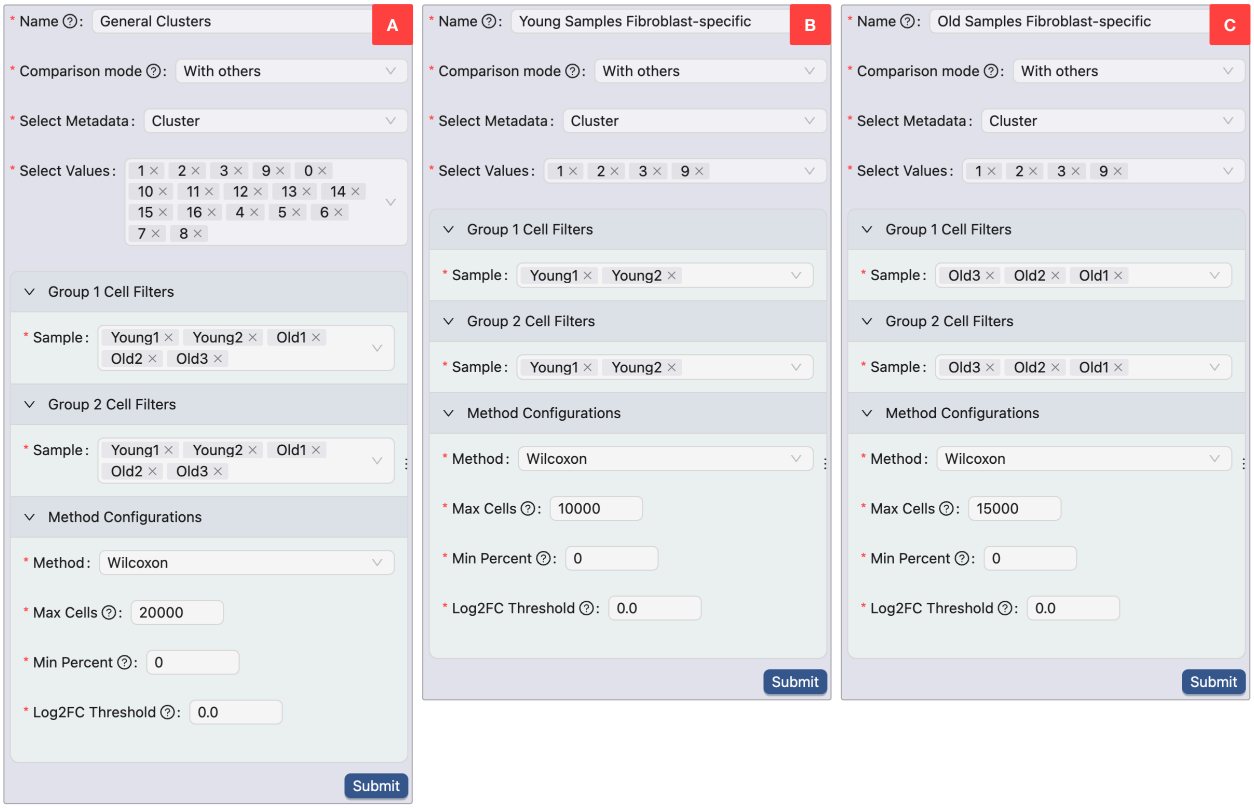

Figure S1: Detailed settings for differential expression analyses in CytoAnalyst. A) Differential expression analysis settings for cluster markers. B) Differential expression analysis settings for young Fibroblast-specific markers. C) Differential expression analysis settings for old Fibroblast-specific markers.

The detailed parameters are as follows:

- *Comparison mode*: "With others"
  Enables comparison of cells from one group against cells from all other groups
- *Select metadata*: "Cluster"
  Specifies cluster information stored in metadata as the grouping variable
- *Select values*: All clusters (1–17)
  For each cluster, compares gene expression of its cells against cells from all other clusters
- *Group 1 Cell Filters*: All five samples
  Includes all cells from the original study's samples in the analysis
- *Group 2 Cell Filters*: All five samples
  Matches Group 1 to ensure comprehensive comparison
- *Statistical test*: Wilcoxon
  Maintains consistency with the original study's statistical methodology
- *Max cells*: 20,000
  Set above the dataset size of 15,457 cells to ensure the inclusion of all cells
- *Min Percent*: 0
  Enables inclusion of all genes regardless of expression percentage

- *Log2FC Threshold*: 0
  Allows detection of all differentially expressed genes regardless of fold change magnitude

Using these settings, we perform 17 differential expression analyses, one for each cluster. Figures S2A– S2I and Figures S3A– S3H present volcano plots for the differential expression analyses of cluster markers, all generated using CytoAnalyst. In each plot, the x-axis displays the log fold change, and the y-axis shows the -log10 of the adjusted p-value. To highlight differentially expressed genes in each comparison, we set thresholds of adjusted p-value $< 0.05$ and average log fold change $> 0.5$. The complete results for all comparisons are provided in Supplementary Table S2 (.xlsx file).

To validate our findings, we compare CytoAnalyst's results with those from the original study. For this comparison, we select genes meeting the following criteria in CytoAnalyst: positive log fold change and expression in at least 25% of cells within the cluster. From these candidates, we select top differentially expressed genes (ranked by lowest p-value) in each cluster, matching the number of genes identified in the original study. Our analysis demonstrates strong concordance with the original study, with CytoAnalyst identifying 88.4% of the genes previously reported. The observed differences can be attributed to our use of Seurat version 5 for differential expression analysis, whereas the original study employed Seurat version 3. Seurat version 5 incorporates improved methods for calculating log fold change and p-value, which likely account for these variations in results.

#### 1.2 DE Analysis to Identify Fibroblast-Specific Markers in Young Samples

In this analysis, we examine clusters 1, 2, 3, and 9, which were identified as Fibroblast subpopulations in the original study, to identify subpopulation-specific markers in young samples. We compare each Fibroblast cluster against all other cells in young samples using CytoAnalyst. The analysis settings are shown in Figure S1B, with the following parameters:

- *Comparison mode*: "With others"
  Enables comparison of cells from one group against cells from all other groups
- *Select metadata*: "Cluster"
  Uses cluster information from metadata as the grouping variable
- *Select values*: Clusters 1, 2, 3, and 9
  Compares each Fibroblast cluster against all other clusters individually (e.g., cluster 1 versus cluster 0 and clusters 2–16)
- *Group 1 Cell Filters*: Young samples
  Restricts analysis to young samples for Fibroblast subpopulation-specific marker identification
- *Group 2 Cell Filters*: Young samples
  Maintains consistent sample selection with Group 1
- *Statistical test*: Wilcoxon
  Maintains consistency with the original study's statistical methodology
- *Max cells*: 10,000
  Set above the young sample size of 5,454 cells to ensure the inclusion of all cells
- *Min Percent*: 0
  Enables inclusion of all genes regardless of expression percentage
- *Log2FC Threshold*: 0
  Allows detection of all differentially expressed genes regardless of fold change magnitude

Using these parameters, we perform four differential expression analyses, one for each Fibroblast subpopulation in young samples. Figure S1B presents the volcano plots for these analyses, generated using CytoAnalyst. Each plot displays log fold change on the x-axis and -log10 of adjusted p-value on the y-axis. To identify significantly differentially expressed genes, we apply thresholds of adjusted p-value $< 0.05$ and

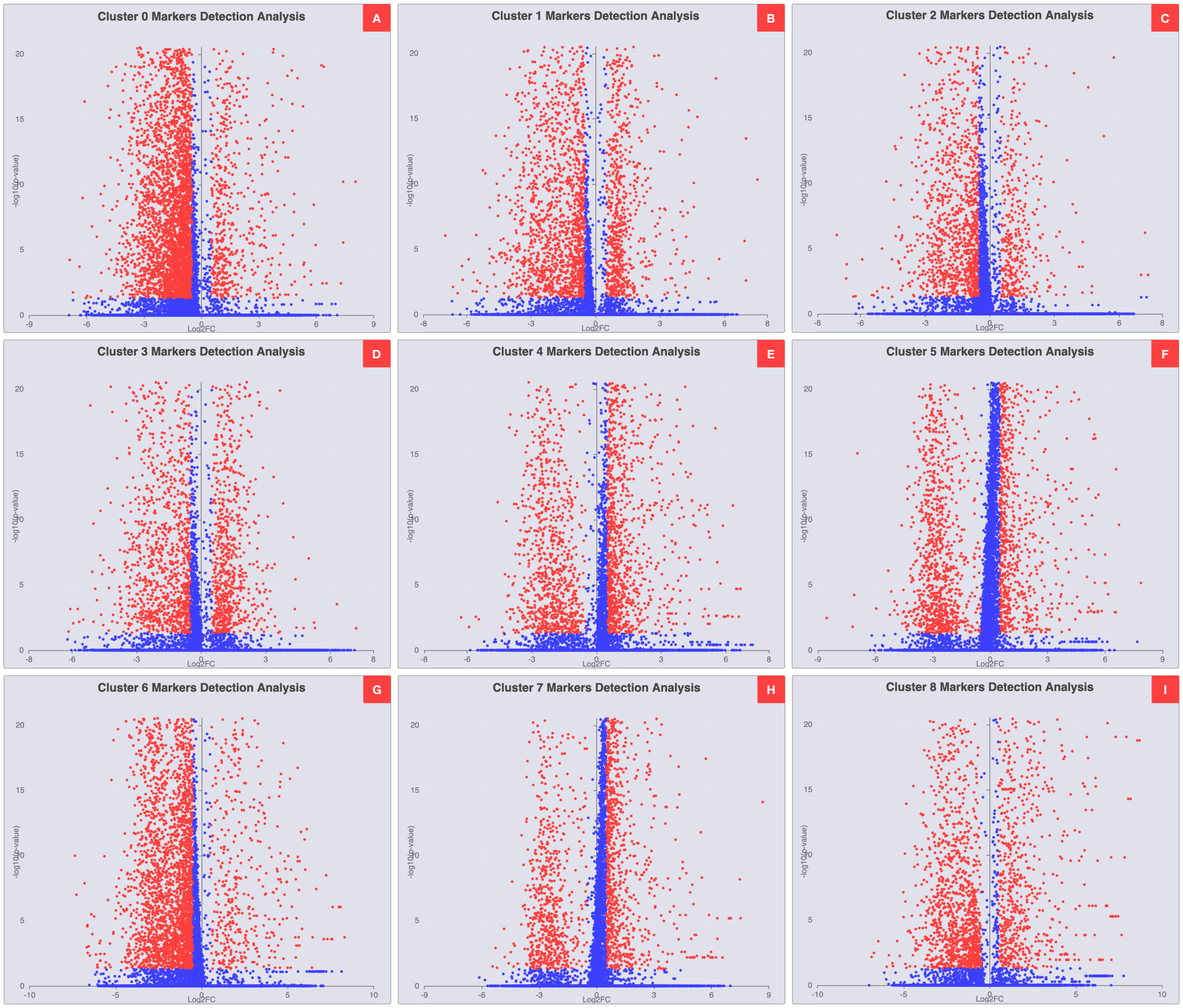

Figure S2: Volcano plots for differential expression analysis of clusters 0–8, The x-axis represents the log fold change, and the y-axis represents the -log10 of the adjusted p-value. The significance threshold is set to adjusted p-value $< 0.05$, and average log fold change $> 0.5$. A) DE analysis results of cluster 0 vs. all other clusters. B) DE analysis results of cluster 1 vs. all other clusters. C) DE analysis results of cluster 2 vs. all other clusters. D) DE analysis results of cluster 3 vs. all other clusters. E) DE analysis results of cluster 4 vs. all other clusters. F) DE analysis results of cluster 5 vs. all other clusters. G) DE analysis results of cluster 6 vs. all other clusters. H) DE analysis results of cluster 7 vs. all other clusters. I) DE analysis results of cluster 8 vs. all other clusters.

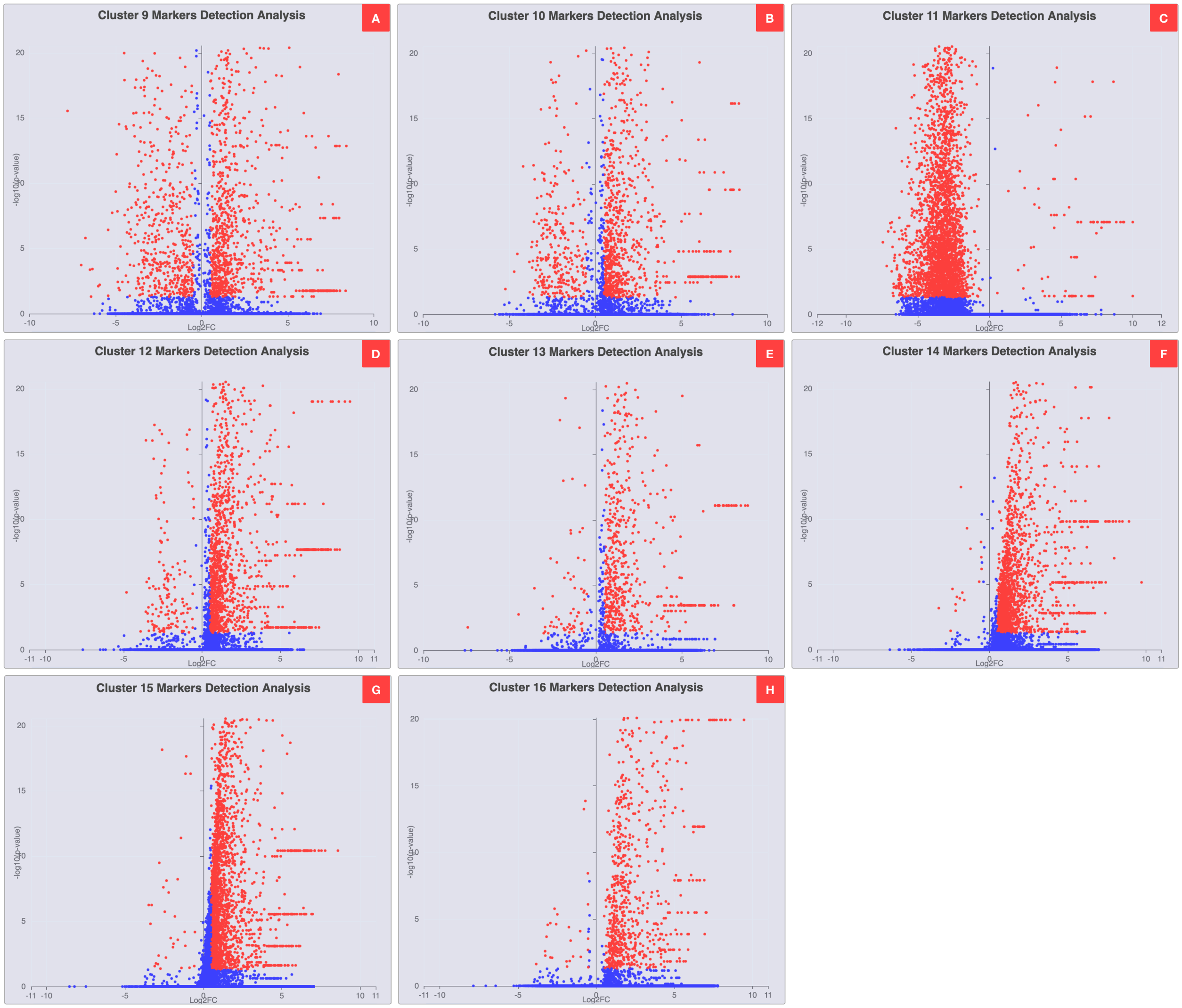

Figure S3: Volcano plots for differential expression analysis of clusters 9–16, The x-axis represents the log fold change, and the y-axis represents the -log10 of the adjusted p-value. The significance threshold is set to adjusted p-value $< 0.05$, and average log fold change $> 0.5$. A) DE analysis results of cluster 9 vs. all other clusters. B) DE analysis results of cluster 10 vs. all other clusters. C) DE analysis results of cluster 11 vs. all other clusters. D) DE analysis results of cluster 12 vs. all other clusters. E) DE analysis results of cluster 13 vs. all other clusters. F) DE analysis results of cluster 14 vs. all other clusters. G) DE analysis results of cluster 15 vs. all other clusters. H) DE analysis results of cluster 16 vs. all other clusters.

average log fold change $> 0.5$. Complete results for all comparisons are available in Supplementary Table S3 (.xlsx file).

Following the same comparison methodology used for general cluster markers, we validate our results against the original study. The analysis demonstrates strong concordance, with CytoAnalyst identifying 91.9% of the genes reported in the original study.

#### 1.3 DE Analysis to Identify Fibroblast-Specific Markers in Old Samples

In this analysis, we analyze clusters 1, 2, 3, and 9 by comparing each of them to other cells in old samples to identify Fibroblast subpopulation-specific markers in old samples. The analysis settings are shown in Figure S1C, with the following parameters:

- *Comparison mode*: "With others"
  Enables comparison of cells from one group against cells from all other groups
- *Select metadata*: "Cluster"
  Uses cluster information from metadata as the grouping variable
- *Select values*: Clusters 1, 2, 3, and 9
  Compares each Fibroblast cluster against all other clusters individually
- *Group 1 Cell Filters*: Old samples
  Restricts analysis to old samples for Fibroblast subpopulation-specific marker identification
- *Group 2 Cell Filters*: Old samples
  Maintains consistent sample selection with Group 1
- *Statistical test*: Wilcoxon
  Maintains consistency with the original study's statistical methodology
- *Max cells*: 15,000
  Set above the old sample size of 10,003 cells to ensure inclusion of all cells
- *Min Percent*: 0
  Enables inclusion of all genes regardless of expression percentage
- *Log2FC Threshold*: 0
  Allows detection of all differentially expressed genes regardless of fold change magnitude

Using these parameters, we perform four differential expression analyses, one for each Fibroblast subpopulation in old samples. Figure S5A–D presents the volcano plots for these analyses, generated using CytoAnalyst. Each plot displays log fold change on the x-axis and -log10 of adjusted p-value on the y-axis. To identify significantly differentially expressed genes, we apply thresholds of adjusted p-value $< 0.05$ and average log fold change $> 0.5$. Complete results for all comparisons are available in Supplementary Table S4 (.xlsx file).

Following the same comparison methodology used for general cluster markers, we validate our results against the original study. The analysis demonstrates strong concordance, with CytoAnalyst identifying 91.1% of the genes reported in the original study.

#### 1.4 DE Analysis Comparing Fibroblast Subtypes in Young Samples

Using the cell type annotations from the original study, we perform an analysis of young sample Fibroblast populations, comprising Secretory-papillary Fibroblasts, Pro-inflammatory Fibroblasts, Secretory-reticular Fibroblasts, and Mesenchymal Fibroblasts. For each Fibroblast subpopulation, we compare its gene expression against all other Fibroblast subpopulations. The analysis settings are shown in Figure S6A–D, with the following parameters:

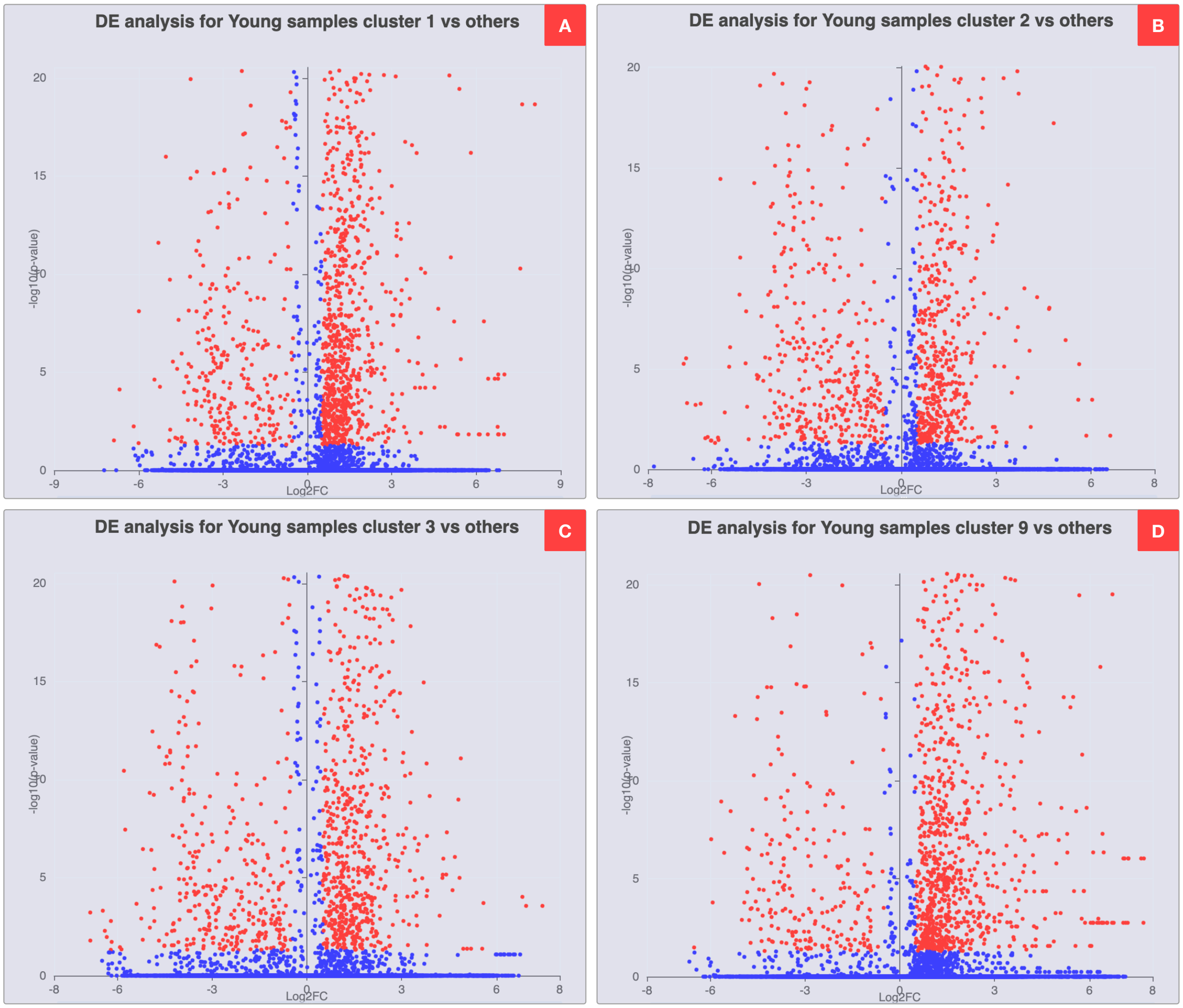

Figure S4: Volcano plots of differential expression analysis of Fibroblast subpopulation-specific markers in young samples. The x-axis represents the log fold change, and the y-axis represents the -log10 of the adjusted p-value. The significance threshold is set to adjusted p-value $< 0.05$, and average log fold change $> 0.5$. A) DE analysis results of cluster 1 vs. other clusters of young samples. B) DE analysis results of cluster 2 vs. other clusters of young samples. C) DE analysis results of cluster 3 vs. other clusters of young samples. D) DE analysis results of cluster 9 vs. other clusters of young samples.

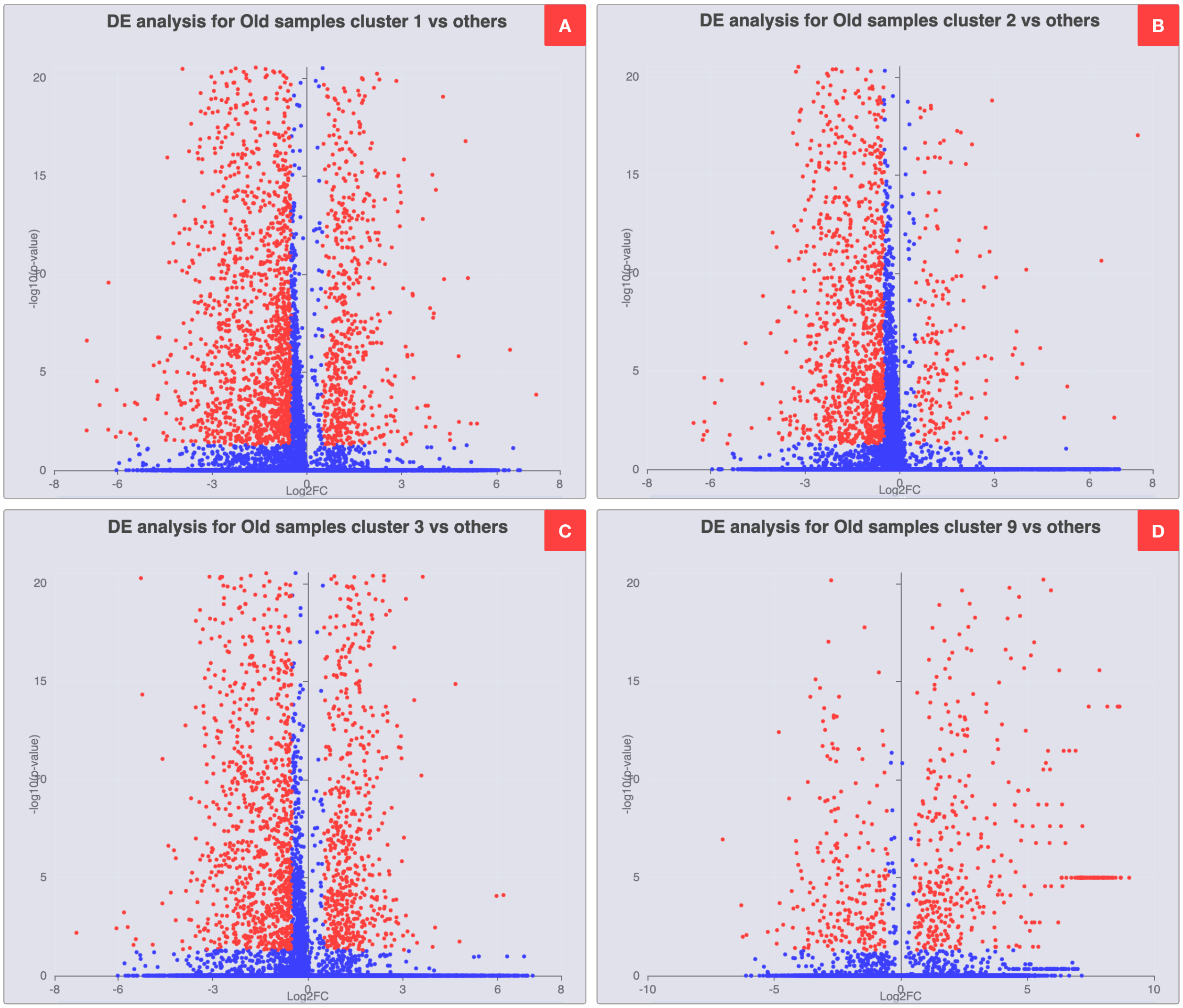

Figure S5: Volcano plots of differential expression analysis of Fibroblast subpopulation-specific markers in old samples. The x-axis represents the log fold change, and the y-axis represents the -log10 of the adjusted p-value. The significance threshold is set to adjusted p-value $< 0.05$, and average log fold change $> 0.5$. A) DE analysis results of cluster 1 vs. all other clusters of old samples. B) DE analysis results of cluster 2 vs. all other clusters of old samples. C) DE analysis results of cluster 3 vs. all other clusters of old samples. D) DE analysis results of cluster 9 vs. all other clusters of old samples.

- *Group 1 Cell Filters*: Young samples
  Restricts analysis to young samples for Fibroblast subpopulation comparison
- *Group 1 Metadata Filters*: "Celltype"
  Uses cell type annotations from metadata for Group 1 classification
- *Group 1 Metadata Values*: Target Fibroblast subpopulation
  Selects specific Fibroblast subpopulation for comparison against others
- *Group 2 Cell Filters*: Young samples
  Maintains consistent sample selection with Group 1
- *Group 2 Metadata Filters*: "Celltype"
  Uses same annotation category as Group 1
- *Group 2 Metadata Values*: All other Fibroblast subpopulations
  Includes remaining Fibroblast subpopulations for comparison
- *Statistical test*: Wilcoxon
  Maintains consistency with the original study's statistical methodology
- *Max cells*: 10,000
  Set above the young Fibroblast sample size of 1,792 cells to ensure inclusion of all cells
- *Min Percent*: 0
  Enables inclusion of all genes regardless of expression percentage
- *Log2FC Threshold*: 0
  Allows detection of all differentially expressed genes regardless of fold change magnitude

Using these parameters, we perform four differential expression analyses, one for each Fibroblast subpopulation in young samples. Figure S7A–D presents volcano plots for these analyses, generated using CytoAnalyst. Each plot displays log fold change on the x-axis and -log10 of adjusted p-value on the y-axis. To identify significantly differentially expressed genes, we apply thresholds of adjusted p-value $< 0.05$ and average log fold change $> 0.5$. Complete results for all comparisons are available in Supplementary Table S5 (.xlsx file).

Following the same comparison methodology used for general cluster markers, we validate our results against the original study. The analysis shows substantial concordance, with CytoAnalyst identifying 84.7% of the genes reported in the original study.

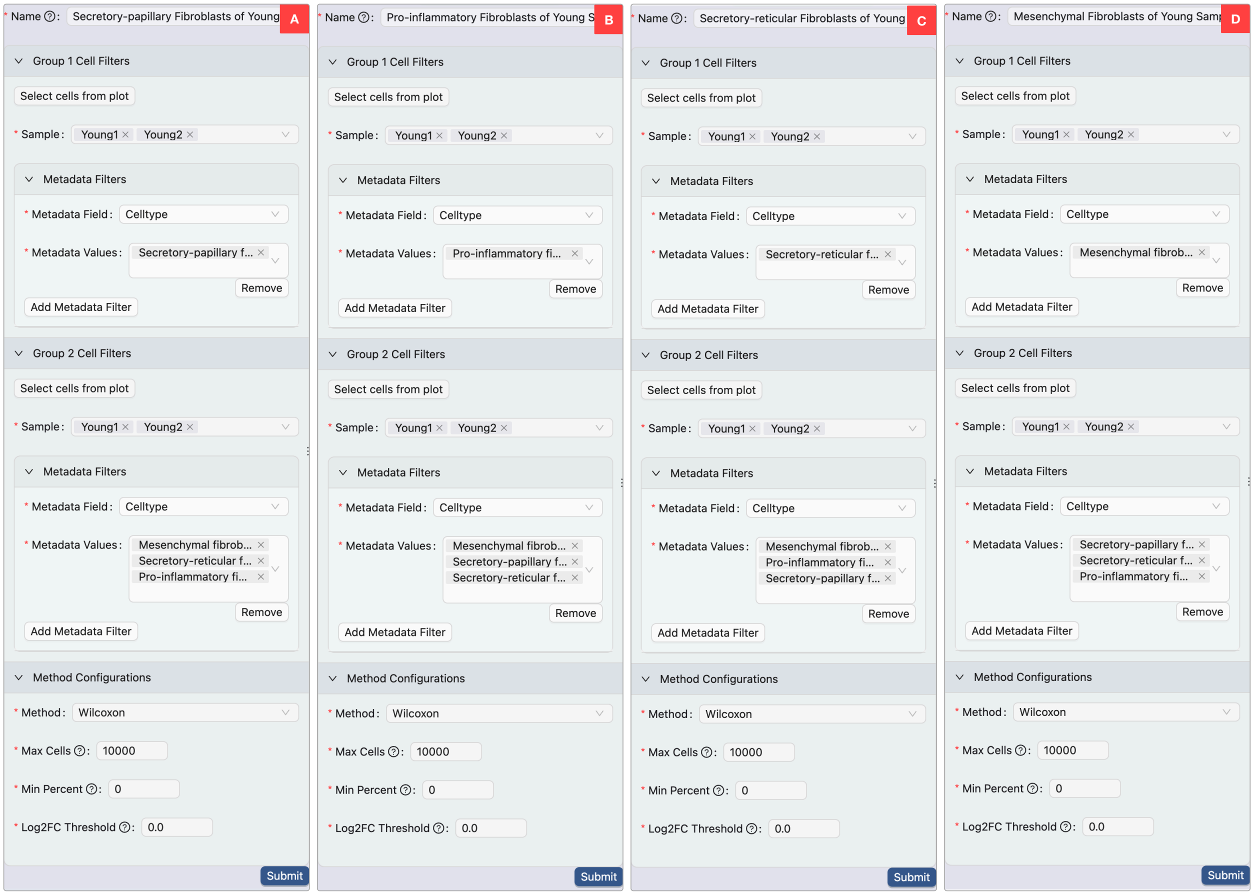

Figure S6: Differential expression analysis settings for Fibroblast subpopulations in young samples. A) DE analysis settings for Secretory-papillary Fibroblasts. B) DE analysis settings for Pro-inflammatory Fibroblasts. C) DE analysis settings for Secretory-reticular Fibroblasts. D) DE analysis settings for Mesenchymal Fibroblasts.

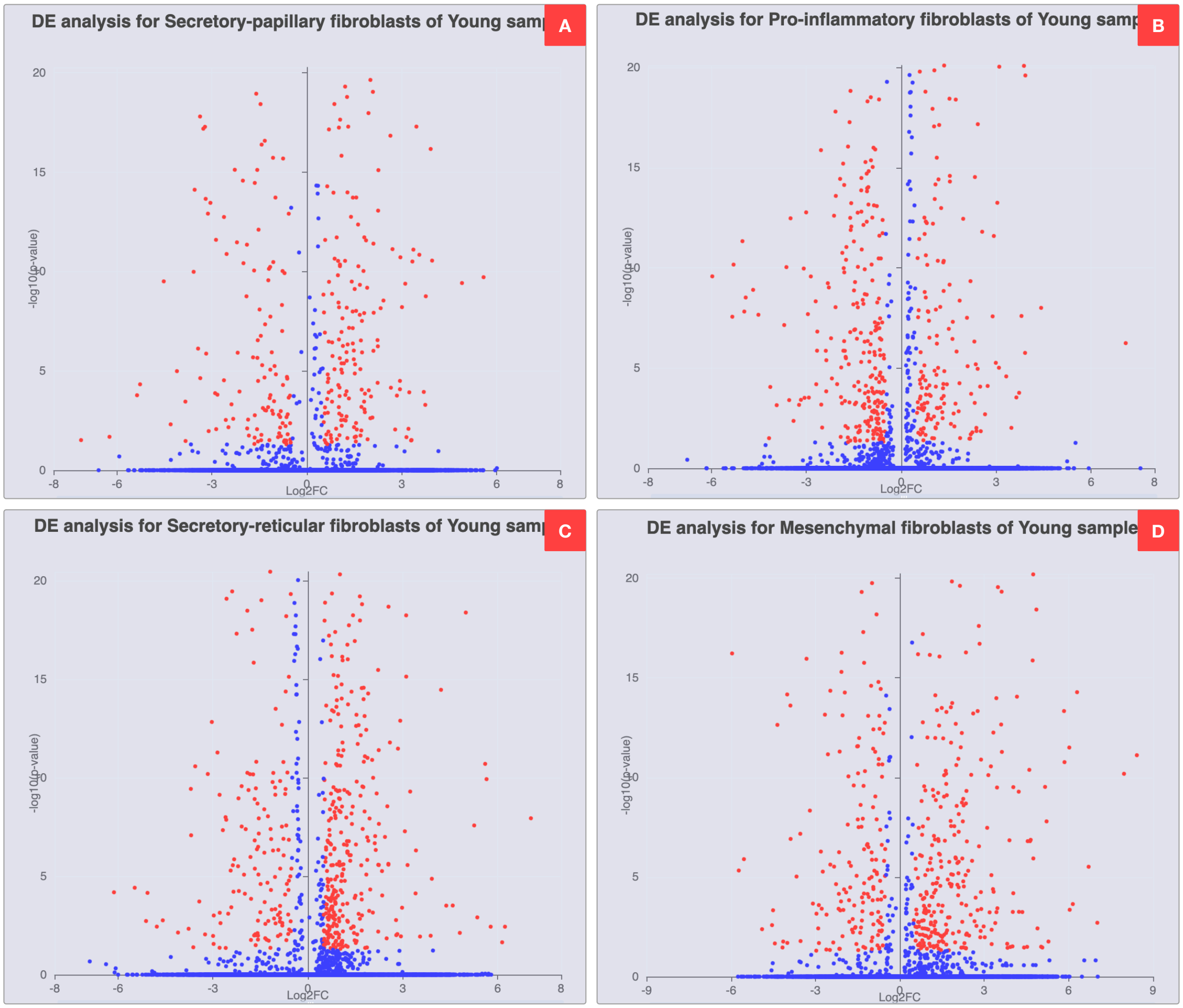

Figure S7: Volcano plots of differential expression analysis of Fibroblast subpopulation-specific markers in young samples. The x-axis represents the log fold change, and the y-axis represents the -log10 of the adjusted p-value. The significance threshold is set to adjusted p-value $< 0.05$, and average log fold change $> 0.5$. A) DE analysis results of Secretory-papillary Fibroblasts vs. all other Fibroblast subpopulations. B) DE analysis results of Pro-inflammatory Fibroblasts vs. all other Fibroblast subpopulations. C) DE analysis results of Secretory-reticular Fibroblasts vs. all other Fibroblast subpopulations. D) DE analysis results of Mesenchymal Fibroblasts vs. all other Fibroblast subpopulations.

#### 2 Case Study 3: Pseudo-Time Trajectory Inference

In this case study, we perform trajectory inference analyses on a bone marrow dataset integrated from three different studies by Dahlin et al. [2], Schaum et al. [3], and Han et al. [4].

In the first study by Dahlin et al., the authors isolated bone marrow cells from C57BL/6 (wild-type) and W41/W41 (c-Kit mutant) mice, aiming to create a comprehensive transcriptional landscape of hematopoietic stem and progenitor cells. The authors used the 10x Genomics Cell Ranger v1.3 pipeline to process the raw sequencing data. For quality control of single-cell data, the authors removed cells with fewer than 500 detected genes, with more than 10% of reads mapping to mitochondrial genes, or with a total number of UMI counts more than 3 times the standard deviation above the mean count. This resulted in a final dataset of 44,802 WT cells and 13,815 W41/W41 cells.

In the second study by The Tabula Muris Consortium, the authors collected single-cell transcriptomic data from 20 mouse organs from three female and four male C57BL/6JN mice aged 10-15 weeks. The study aimed at creating a comprehensive single-cell atlas across mouse tissues. The authors captured cells using two different methods: fluorescence-activated-cell-sorting-based (FACS-based) capture in plates and microfluidic-droplet-based capture. The authors used STAR version 2.5.2b and HTSEQ version 0.6.1p1 to align sequences from the FACS-based method and obtain the gene count data, respectively. For the microfluidic-droplet-based method, the authors used Cell Ranger version 2.0.1 for alignment and gene counts. For quality control, the authors excluded cells with fewer than 500 detected genes, fewer than 50,000 reads (FACS-based), or fewer than 1,000 UMIs (microfluidic-droplet-based). This resulted in a final dataset of 44,949 cells from the FACS-based method and 55,656 cells from the microfluidic-droplet-based method.

In the third study by Han et al., the authors developed Microwell-seq, a high-throughput and cost-effective method for single-cell RNA sequencing, to construct a comprehensive Mouse Cell Atlas (MCA). They collected tissue samples from 6-10 week-old adult C57BL/6 mice, E14.5 embryos, and 1-day-old neonatal mice, spanning more than 50 different mouse tissues, organs, and cell cultures. The authors used STAR version 2.5.2a to align the raw sequencing data and obtain the gene counts. For quality control, the authors excluded cells with fewer than 500 expressed transcripts and cells with mitochondrial gene percentages exceeding 10%. To mitigate batch effects in cross-tissue comparisons, they removed batch-specific background gene expression, defined as the average gene detection for cellular barcodes with less than 500 UMI, multiplied by a coefficient of 2. This resulted in a final dataset of more than 400,000 cells across all tissues.

In this case study, we retrieve the integrated dataset from the three studies processed by Björklund et al. (`https://nbisweden.github.io/workshop-scRNAseq/labs/seurat/seurat_07_trajectory.html`). The dataset contains 5,828 cells and has been preprocessed using the Seurat package, including cell filtering to retain only cells in the bone marrow tissue, normalization, feature selection, dimensionality reduction, and clustering. In addition, Björklund et al. also removed clusters that are disconnected or outlier cells. Using this dataset, Björklund et al. performed trajectory inference analyses using the Slingshot method to identify developmental trajectories of bone marrow cells, focusing on B cell lineage, Granulocyte lineage, Dendritic cell lineage, HSC progenitor, T cell lineage, Monocyte lineage, Mast Cell lineage, RBC lineage, and Megakaryocyte cell lineage. Finally, the authors used the tradeSeq method to identify genes that change during the developmental trajectories of each lineage. Here, we aim to reproduce the trajectory inference results from the time trajectory analysis workshop by Björklund et al. using our CytoAnalyst platform, focusing on the B cell lineage, Granulocyte lineage, Dendritic cell lineage, and HSC progenitor.

Using the available embeddings and complete clustering results from the dataset metadata, we perform trajectory inference on four lineage-specific markers (Table S1): Ms4a1 (B cell lineage), Ltf (Granulocyte lineage), Siglech (Dendritic cell lineage), and Cd34 (HSC progenitor). These cell-type markers were identified in the original analysis.

Table S1: A list of cell-type labels corresponding to each marker

| Cell Type | Marker |
|---|---|
| B cell lineage | Ms4a1 |
| HSPC lineage | Cd34 |
| Granulocyte lineage | Ltf |
| Dendritic cell lineage | Siglech |

We perform four distinct trajectory inference analyses using the Slingshot method, one for each lineage-specific gene.

#### 2.1 Trajectory Inference for the Ms4a1 Gene

Figure S8 presents the clustering results from the authors, which have been refined by removing disconnected and outlier clusters alongside Ms4a1 gene expression patterns. The analysis reveals high Ms4a1 expression in cluster 20 and minimal expression in clusters 33 and 47. Based on these expression patterns, we designate cluster 20 as the starting point and clusters 33 and 47 as endpoints for the Slingshot trajectory inference parameters.

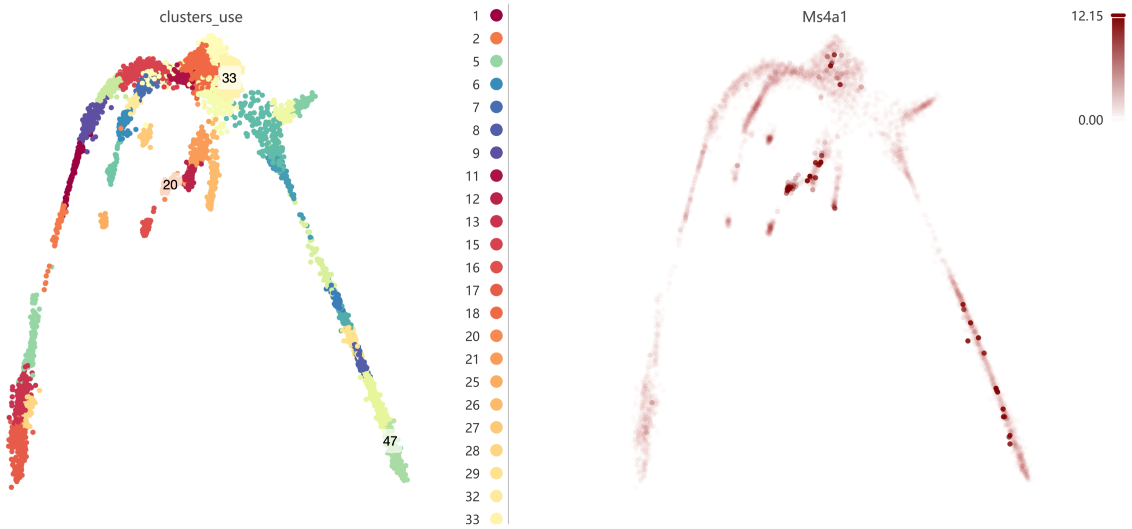

Figure S8: Clustering result (left) and Ms4a1 gene expression (right).

We then perform trajectory inference on the Ms4a1 gene using CytoAnalyst. The analysis settings are shown in Figure S9A, with the following parameters:

- *Name*: Analysis identifier
- *Method*: "Slingshot"
  Trajectory inference method used for analysis
- *Embeddings*: "umap3d"
  UMAP-generated embedding from the original study
- *Group cells by*: "Metadata"
  Specifies cell grouping method
- *Metadata*: "cluster_use"
  Metadata field for cell grouping
- *Start Groups*: "20"
  Initial cluster for trajectory analysis
- *End Groups*: "33", "47"
  Terminal clusters for trajectory analysis
- *Distance method*: "Mutual nearest neighbors"
  Method for computing inter-cluster distances

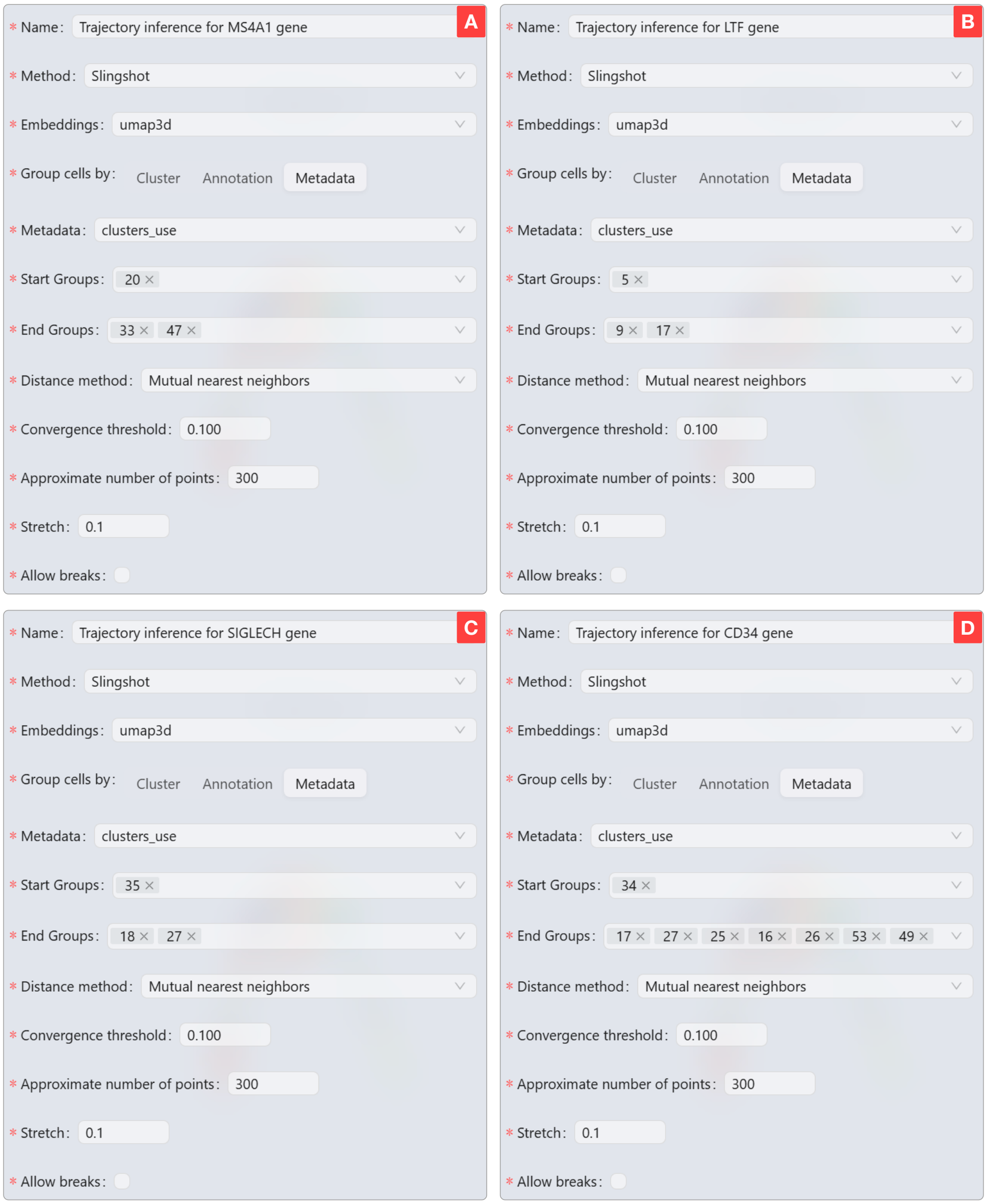

Figure S9: Parameters used for trajectory inference analyses in CytoAnalyst. (1) The trajectory inference settings for the Ms4a1 gene. (2) The trajectory inference settings for the Ltf gene. (3) The trajectory inference settings for the Siglech gene. (4) The trajectory inference settings for the Cd34 gene.

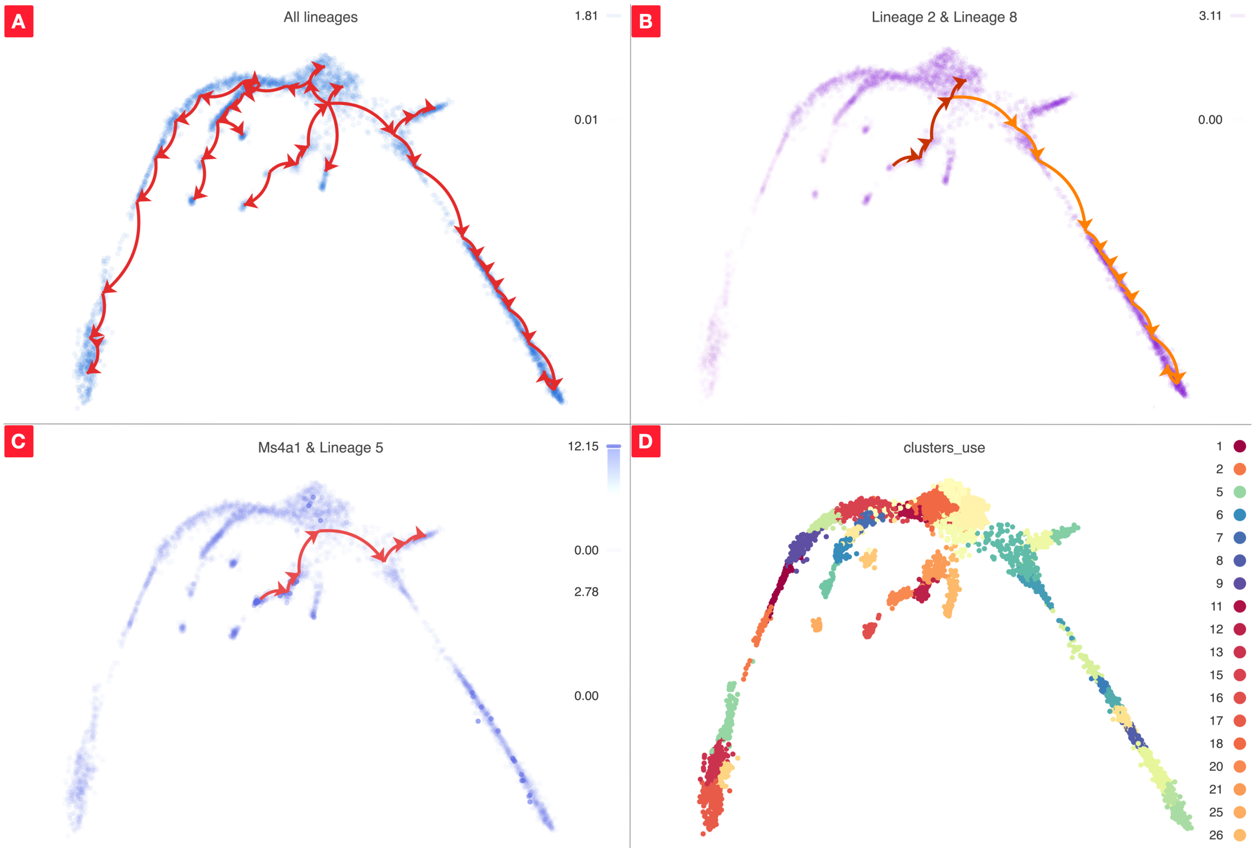

Figure S10: Trajectory inference analysis results for the Ms4a1 gene. A) Complete trajectory lineage map. B) Developmental pathways (lineages 2 and 8) from starting to ending groups. C) Ms4a1 gene expression with lineage 5 overlay. D) Original clustering results with highlighted start and end groups.

- *Convergence threshold*: "0.100"
  Threshold for change in total distance between cells and curve projections

- *Approximate number of points*: "300"
  Number of points along trajectory curves

- *Stretch*: "0.1"
  Curve extrapolation factor beyond endpoints

- *Allow breaks*: "False"
  Ensures continuous principal curves from origin

Figure S10 presents the trajectory inference analysis results for the Ms4a1 gene:

- Panel A displays all identified trajectory lineages
- Panel B shows developmental pathways from the starting to ending groups
- Panel C overlays lineage 5 with Ms4a1 gene expression, illustrating expression changes along the trajectory
- Panel D presents the original study's clustering results, which guided the selection of start and end groups based on expression patterns

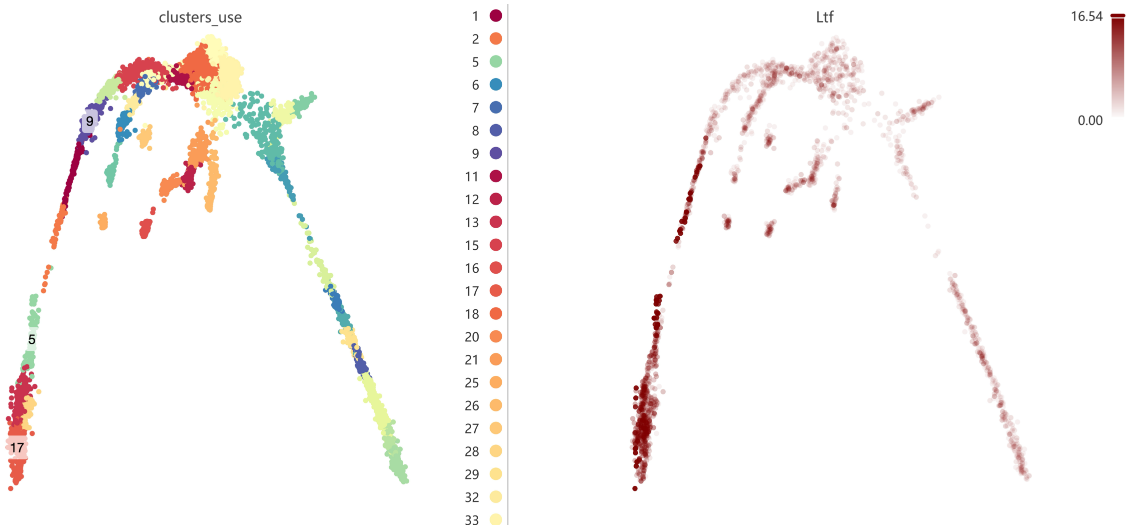

Figure S11: Clustering result (left) and Ltf gene expression (right).

#### 2.2 Trajectory Inference for the Ltf, Siglech, and Cd34 Genes

We perform trajectory inference analyses for Ltf, Siglech, and Cd34 genes following the methodology established for Ms4a1. Using clustering results and gene expression patterns, we identify optimal starting and ending groups for each gene (Figures S11, S12, and S13). Based on expression patterns, we designate:

- Ltf gene: cluster 5 (start), clusters 9 and 17 (end)
- Siglech gene: cluster 35 (start), clusters 18 and 27 (end)
- Cd34 gene: cluster 34 (start), clusters 17, 27, 25, 16, 26, 53, and 49 (end)

We conduct trajectory inference analyses using CytoAnalyst with parameters matching those used for Ms4a1 (detailed settings shown in Figure S9B–D).

Results for each gene are presented in Figures S14, S15, and S16, respectively. Each figure comprises four panels:

- Panel A: Complete trajectory lineage map
- Panel B: Developmental pathways from start to end groups
- Panel C: Gene expression overlay with representative lineage
- Panel D: Original clustering results with highlighted groups

#### 2.3 Comparison with Original Study Results

The original study conducted trajectory inference analysis for the Cd34 gene and provided a lineage table for their results. We compare our CytoAnalyst-derived results with the original analysis, as shown in Figure S17. The lineage tables demonstrate perfect concordance between the two analyses.

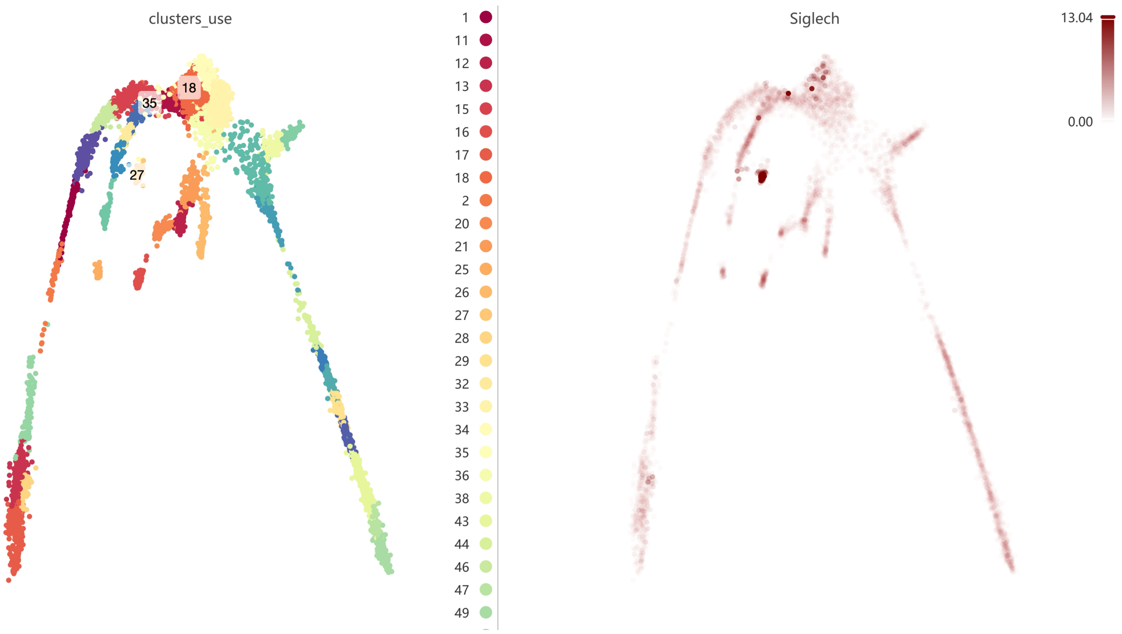

Figure S12: Clustering result (left) and Siglech gene expression (right).

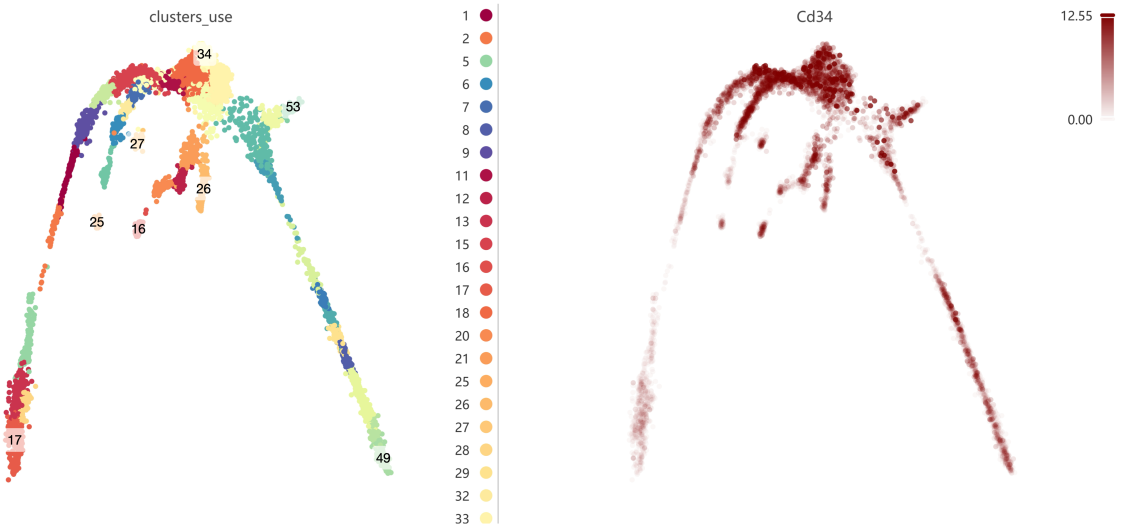

Figure S13: Clustering result (left) and Cd34 gene expression (right).

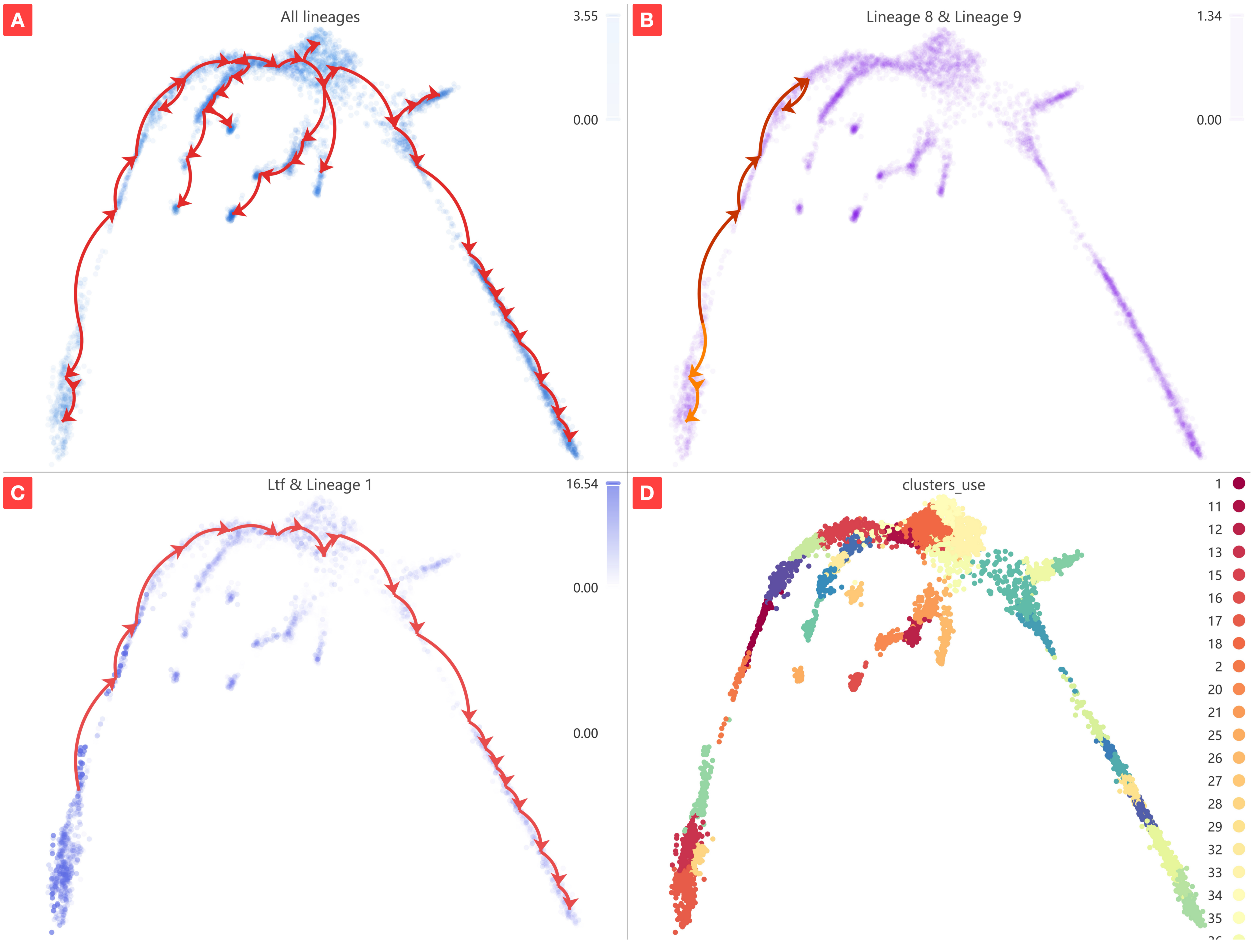

Figure S14: Trajectory inference analysis results for the Ltf gene. A) Complete trajectory lineage map. B) Developmental pathways (lineages 8 and 9) from starting to ending groups. C) Ltf gene expression with lineage 1 overlay. D) Original clustering results with highlighted start and end groups.

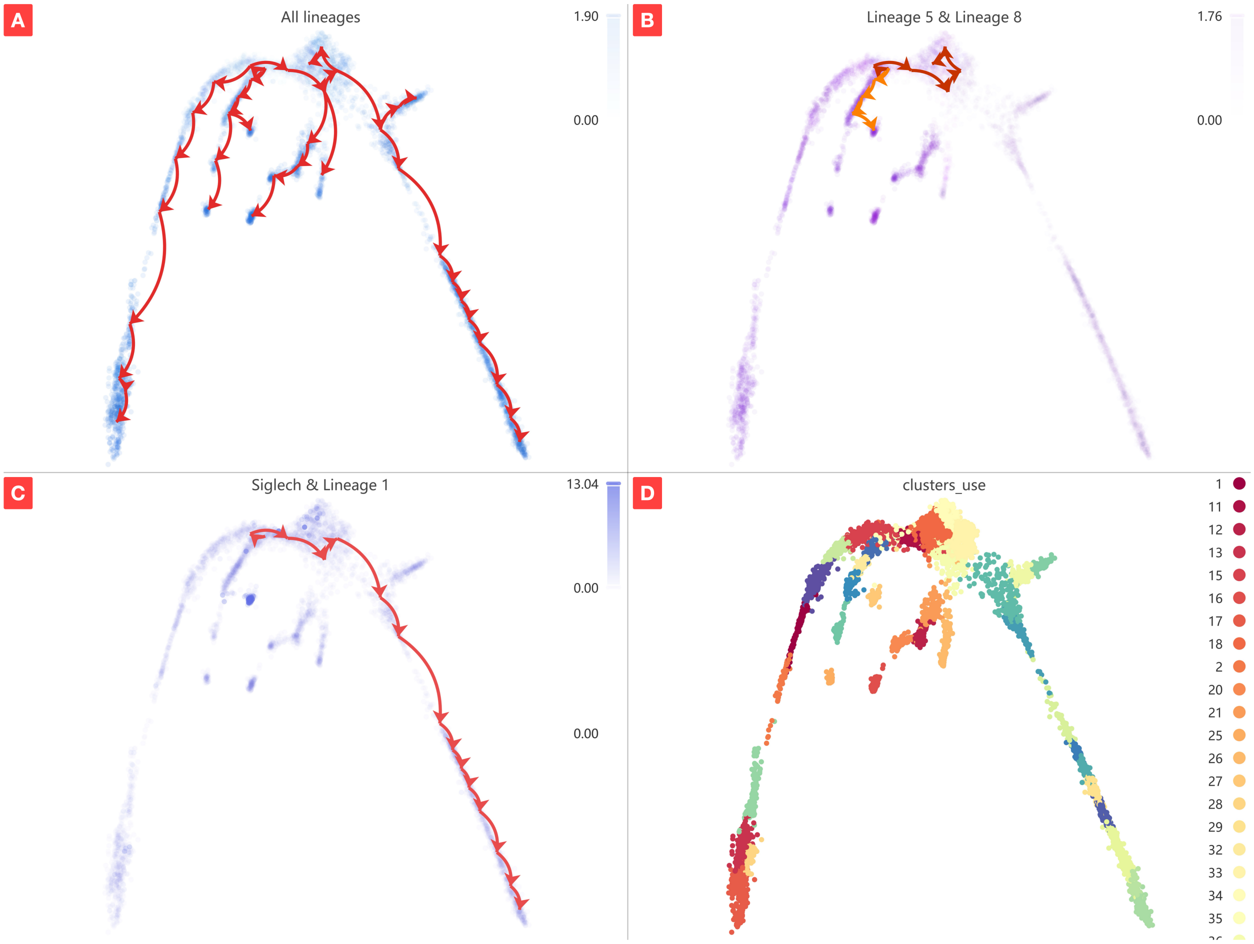

Figure S15: Trajectory inference analysis results for the Siglech gene. A) Complete trajectory lineage map. B) Developmental pathways (lineages 5 and 8) from starting to ending groups. C) Siglech gene expression with lineage 1 overlay. D) Original clustering results with highlighted start and end groups.

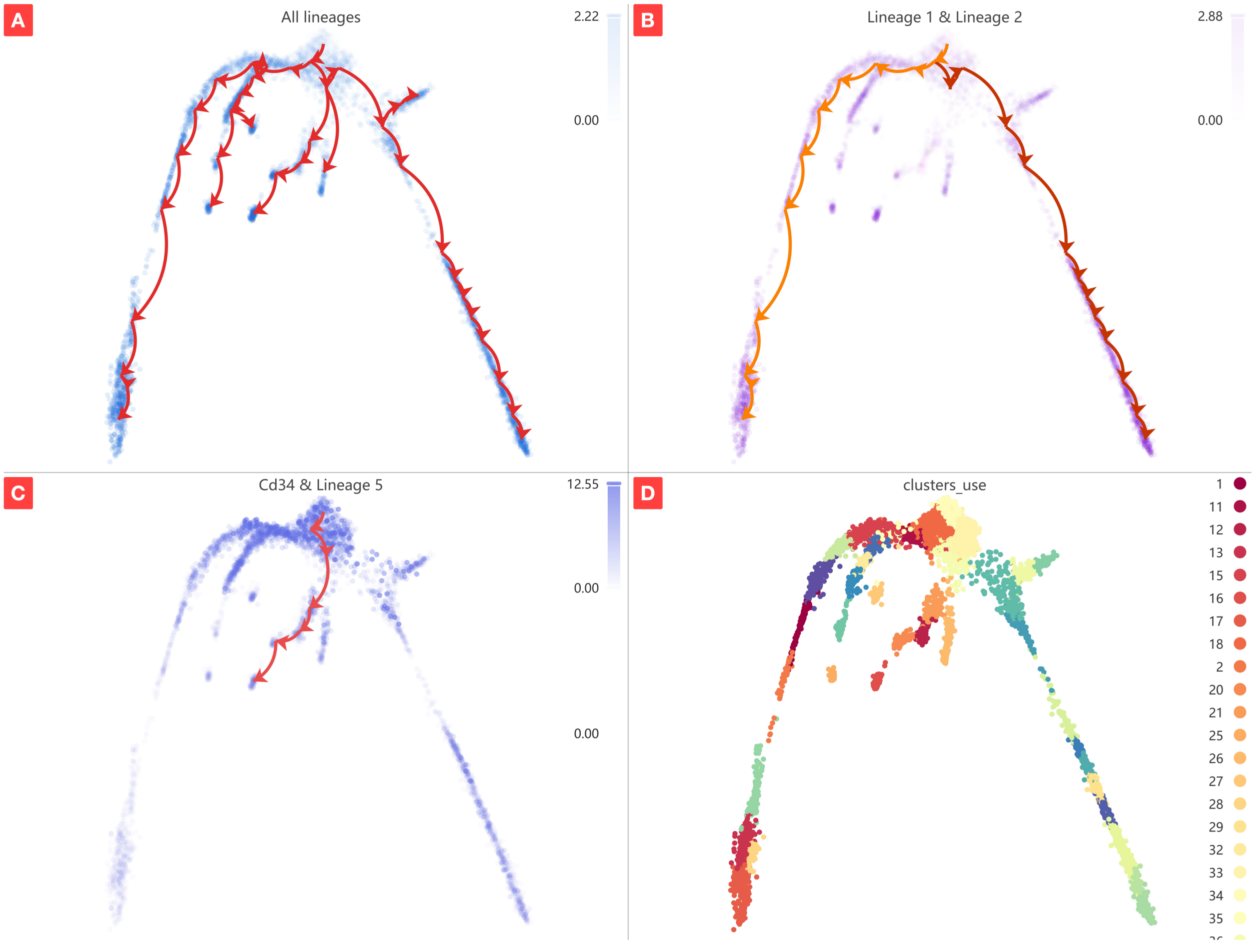

Figure S16: Trajectory inference analysis results for the Cd34 gene. A) Complete trajectory lineage map. B) Developmental pathways (lineages 1 and 2) from starting to ending groups (17 and 49). C) Cd34 gene expression with lineage 5 overlay. D) Original clustering results with highlighted start and end groups.

A

| # | Lineage |
|---|---|
| 1 | 34 → 18 → 36 → 33 → 55 → 59 → 44 → 60 → 58 → 29 → 8 → 43 → 47 → 49 |
| 2 | 34 → 18 → 11 → 15 → 46 → 9 → 1 → 2 → 5 → 13 → 28 → 17 |
| 3 | 34 → 18 → 11 → 15 → 35 → 7 → 32 → 6 → 54 → 25 |
| 4 | 34 → 18 → 11 → 15 → 35 → 7 → 32 → 6 → 27 |
| 5 | 34 → 18 → 36 → 21 → 12 → 20 → 16 |
| 6 | 34 → 18 → 36 → 33 → 55 → 38 → 53 |
| 7 | 34 → 18 → 36 → 26 |

B

```
lineages: 7
Lineage1: 34  18  36  33  55  59  44  60  58  29  8  43  47  49
Lineage2: 34  18  11  15  46  9  1  2  5  13  28  17
Lineage3: 34  18  11  15  35  7  32  6  54  25
Lineage4: 34  18  11  15  35  7  32  6  27
Lineage5: 34  18  36  21  12  20  16
Lineage6: 34  18  36  33  55  38  53
Lineage7: 34  18  36  26
```

Figure S17: Comparison of lineage results from CytoAnalyst and original study. A) CytoAnalyst-derived lineages. B) Original study lineages.

### References

[1] Llorenç Solé-Boldo, Günter Raddatz, Sabrina Schütz, Jan-Philipp Mallm, Karsten Rippe, Anke S Lonsdorf, Manuel Rodríguez-Paredes, and Frank Lyko. Single-cell transcriptomes of the human skin reveal age-related loss of fibroblast priming. *Communications Biology*, 3(1):188, 2020.

[2] Joakim S. Dahlin, Fiona K. Hamey, Blanca Pijuan-Sala, Mairi Shepherd, Winnie WY Lau, Sonia Nestorowa, Caleb Weinreb, Samuel Wolock, Rebecca Hannah, Evangelia Diamanti, David G. Kent, Berthold Göttgens, and Nicola K. Wilson. A single-cell hematopoietic landscape resolves 8 lineage trajectories and defects in Kit mutant mice. *Blood, The Journal of the American Society of Hematology*, 131(21):e1–e11, 2018.

[3] Nicholas Schaum, Jim Karkanias, Norma F. Neff, Andrew P. May, Stephen R. Quake, Tony Wyss-Coray, Spyros Darmanis, Joshua Batson, Olga Botvinnik, Michelle B. Chen, Steven Chen, Foad Green, Robert C. Jones, Ashley Maynard, Lolita Penland, Angela Oliveira Pisco, Rene V. Sit, Geoffrey M. Stanley, James T. Webber, Fabio Zanini, Ankit S. Baghel, Isaac Bakerman, Ishita Bansal, Daniela Berdnik, Biter Bilen, Douglas Brownfield, Corey Cain, Michelle B. Chen, Steven Chen, Min Cho, Giana Cirolia, Stephanie D. Conley, Spyros Darmanis, Aaron Demers, Kubilay Demir, Antoine de Morree, Tessa Divita, Haley du Bois, Laughing Bear Torrez Dulgeroff, Hamid Ebadi, F. Hernán Espinoza, Matt Fish, Qiang Gan, Benson M. George, Astrid Gillich, Foad Green, Geraldine Genetiano, Xueying Gu, Gunsagar S. Gulati, Yan Hang, Shayan Hosseinzadeh, Albin Huang, Tal Iram, Taichi Isobe, Feather Ives, Robert C. Jones, Kevin S. Kao, Guruswamy Karnam, Aaron M. Kershner, Bernhard M. Kiss, William Kong, Maya E. Kumar, Jonathan Y. Lam, Davis P. Lee, Song E. Lee, Guang Li, Qingyun Li, Ling Liu, Annie Lo, Wan-Jin Lu, Anoop Manjunath, Andrew P. May, Kaia L. May, Oliver L. May, Ashley Maynard, Marina McKay, Ross J. Metzger, Marco Mignardi, Dullei Min, Ahmad N. Nabhan, Norma F. Neff, Katharine M. Ng, Joseph Noh, Rasika Patkar, Weng Chuan Peng, Lolita Penland, Robert Puccinelli, Eric J. Rulifson, Nicholas Schaum, Shaheen S. Sikandar, Rahul Sinha, Rene V. Sit, Krzysztof Szade, Weilun Tan, Cristina Tato, Krissie Tellez, Kyle J. Travaglini, Carolina Tropini, Lucas Waldburger, Linda J. van Weele, Michael N. Wosczyna, Jinyi Xiang, Soso Xue, Justin Youngyunpipatkul, Fabio Zanini, Macy E. Zardeneta, Fan Zhang, Lu Zhou, Ishita Bansal, Steven Chen, Min Cho, Giana Cirolia, Spyros Darmanis, Aaron Demers, Tessa Divita, Hamid Ebadi, Geraldine Genetiano, Foad Green, Shayan Hosseinzadeh, Feather Ives, Annie Lo, Andrew P. May, Ashley Maynard, Marina McKay, Norma F. Neff, Lolita Penland, Rene V. Sit, Weilun Tan, Lucas Waldburger, Justin Youngyunpipatkul, Joshua Batson, Olga Botvinnik, Paola Castro, Derek Croote, Spyros Darmanis, Joseph L. DeRisi, Jim Karkanias, Angela Oliveira Pisco, Geoffrey M. Stanley, James T. Webber, Fabio Zanini, Ankit S. Baghel, Isaac Bakerman, Joshua Batson, Biter Bilen, Olga Botvinnik, Douglas Brownfield, Michelle B. Chen, Spyros Darmanis, Kubilay Demir, Antoine de Morree, Hamid Ebadi, F. Hernán Espinoza, Matt Fish, Qiang Gan, Benson M. George, Astrid Gillich, Xueying Gu, Gunsagar S. Gulati, Yan Hang, Albin Huang, Tal Iram, Taichi Isobe, Guruswamy Karnam, Aaron M. Kershner, Bernhard M. Kiss, William Kong, Christin S. Kuo, Jonathan Y. Lam, Benoit Lehallier, Guang Li, Qingyun Li, Ling Liu, Wan-Jin Lu, Dullei Min, Ahmad N. Nabhan, Katharine M. Ng, Patricia K. Nguyen, Rasika Patkar, Weng Chuan Peng, Lolita Penland, Eric J. Rulifson, Nicholas Schaum, Shaheen S. Sikandar, Rahul Sinha, Krzysztof Szade, Serena Y. Tan, Krissie Tellez, Kyle J. Travaglini, Carolina Tropini, Linda J. van Weele, Bruce M. Wang, Michael N. Wosczyna, Jinyi Xiang, Hanadie Yousef, Lu Zhou, Joshua Batson, Olga Botvinnik, Steven Chen, Spyros Darmanis, Foad Green, Andrew P. May, Ashley Maynard, Angela Oliveira Pisco, Stephen R. Quake, Nicholas Schaum, Geoffrey M. Stanley, James T. Webber, Tony Wyss-Coray, Fabio Zanini, Philip A. Beachy, Charles K. F. Chan, Antoine de Morree, Benson M. George, Gunsagar S. Gulati, Yan Hang, Kerwyn Casey Huang, Tal Iram, Taichi Isobe, Aaron M. Kershner, Bernhard M. Kiss, William Kong, Guang Li, Qingyun Li, Ling Liu, Wan-Jin Lu, Ahmad N. Nabhan, Katharine M. Ng, Patricia K. Nguyen, Weng Chuan Peng, Eric J. Rulifson, Nicholas Schaum, Shaheen S. Sikandar, Rahul Sinha, Krzysztof Szade, Kyle J. Travaglini, Carolina Tropini, Bruce M. Wang, Kenneth Weinberg, Michael N. Wosczyna, Sean M. Wu, Hanadie Yousef, Ben A. Barres, Philip A. Beachy, Charles K. F. Chan, Michael F. Clarke, Spyros Darmanis, Kerwyn Casey Huang, Jim Karkanias, Seung K. Kim, Mark A. Krasnow, Maya E. Kumar, Christin S. Kuo, Andrew P. May, Ross J. Metzger, Norma F. Neff, Roel Nusse, Patricia K. Nguyen, Thomas A. Rando, Justin Sonnenburg, Bruce M. Wang, Kenneth Weinberg,

Irving L. Weissman, Sean M. Wu, Stephen R. Quake, Tony Wyss-Coray, The Tabula Muris Consortium, Overall coordination, Logistical coordination, Organ collection and processing, Library preparation and sequencing, Computational data analysis, Cell type annotation, Writing group, Supplemental text writing group, and Principal investigators. Single-cell transcriptomics of 20 mouse organs creates a Tabula Muris. *Nature*, 562(7727):367–372, 2018.

[4] Xiaoping Han, Renying Wang, Yincong Zhou, Lijiang Fei, Huiyu Sun, Shujing Lai, Assieh Saadatpour, Ziming Zhou, Haide Chen, Fang Ye, Daosheng Huang, Yang Xu, Wentao Huang, Mengmeng Jiang, Xinyi Jiang, Jie Mao, Yao Chen, Chenyu Lu, Jin Xie, Qun Fang, Yibin Wang, Rui Yue, Tiefeng Li, He Huang, Stuart H. Orkin, Guo-Cheng Yuan, Ming Chen, and Guoji Guo. Mapping the Mouse Cell Atlas by Microwell-Se. *Cell*, 172(5):1091–1107, 2018.
